# dCas9 regulator to neutralize competition in CRISPRi circuits

**DOI:** 10.1101/2020.08.11.246561

**Authors:** Hsin-Ho Huang, Massimo Bellato, Yili Qian, Pablo Cárdenas, Lorenzo Pasotti, Paolo Magni, Domitilla Del Vecchio

## Abstract

CRISPRi-mediated gene repression allows simultaneous control of many genes. However, despite highly specific sgRNA-promoter binding, multiple sgRNAs still interfere with one another by competing for dCas9. We created a dCas9 regulator that adjusts dCas9 concentration based on sgRNAs’ demand, mitigating competition in CRISPRi-based logic gates. The regulator’s performance is demonstrated on both single-stage and layered CRISPRi logic gates and in two common *E. coli* strains. When a competitor sgRNA causes between two and ~25 fold-change in a logic gate’s input/output response without dCas9 regulator, the response is essentially unchanged when the regulator is used. The dCas9 regulator thus enables concurrent and independent operation of multiple sgRNAs, thereby supporting independent control of multiple genes.

---

The clustered regularly interspaced short palindromic repeats (CRISPR)–dCas9 system allows to create many orthogonal transcriptional regulators that can be used concurrently to manipulate the transcriptome [1, 2] and to engineer layered logic gates for sophisticated computations [3, 4]. Catalytically inactive Cas9 (dCas9) is recruited by a single guide RNA (sgRNA) to a desired target sequence within a promoter to block RNA polymerase (RNAP), a process known as CRISPR interference (CRISPRi) [5]. Hence, the dCas9-sgRNA complex effectively functions as a transcriptional repressor. Because it is possible to change the promoter target by only modifying the sgRNA and the sgRNA-promoter binding is highly specific, one can create a large library of orthogonal transcriptional repressors that, in principle, do not interfere with one another [6]. Because of the high specificity of sgRNA-promoter binding and the lower loading to gene expression resources than with protein-based transcription factors, the CRISPRi-dCas9 system appears as a promising solution to create increasingly large and sophisticated transcriptional programs [7].

Despite the high specificity of sgRNA-promoter binding, multiple sgRNAs can still interfere with one another by competing for limited dCas9 [8–11]. Specifically, Zhang et al. showed that the fold-repression exerted by any one sgRNA through CRISPRi can decrease by up to 5 times when additional sgRNAs are expressed in the circuit [8]. In [9], the authors observed that the ability of one sgRNA to repress its target was hampered when a second sgRNA without a target was constitutively expressed, consistent with predictions from mathematical models [10]. When a new sgRNA is expressed and binds to dCas9, this protein is sequestered away from other sgRNAs, thereby reducing the available dCas9 that can bind to these sgRNAs (apo-dCas9). That is, sgRNAs effectively “load” apo-dCas9 (Figure 1a).

**Figure 1:**
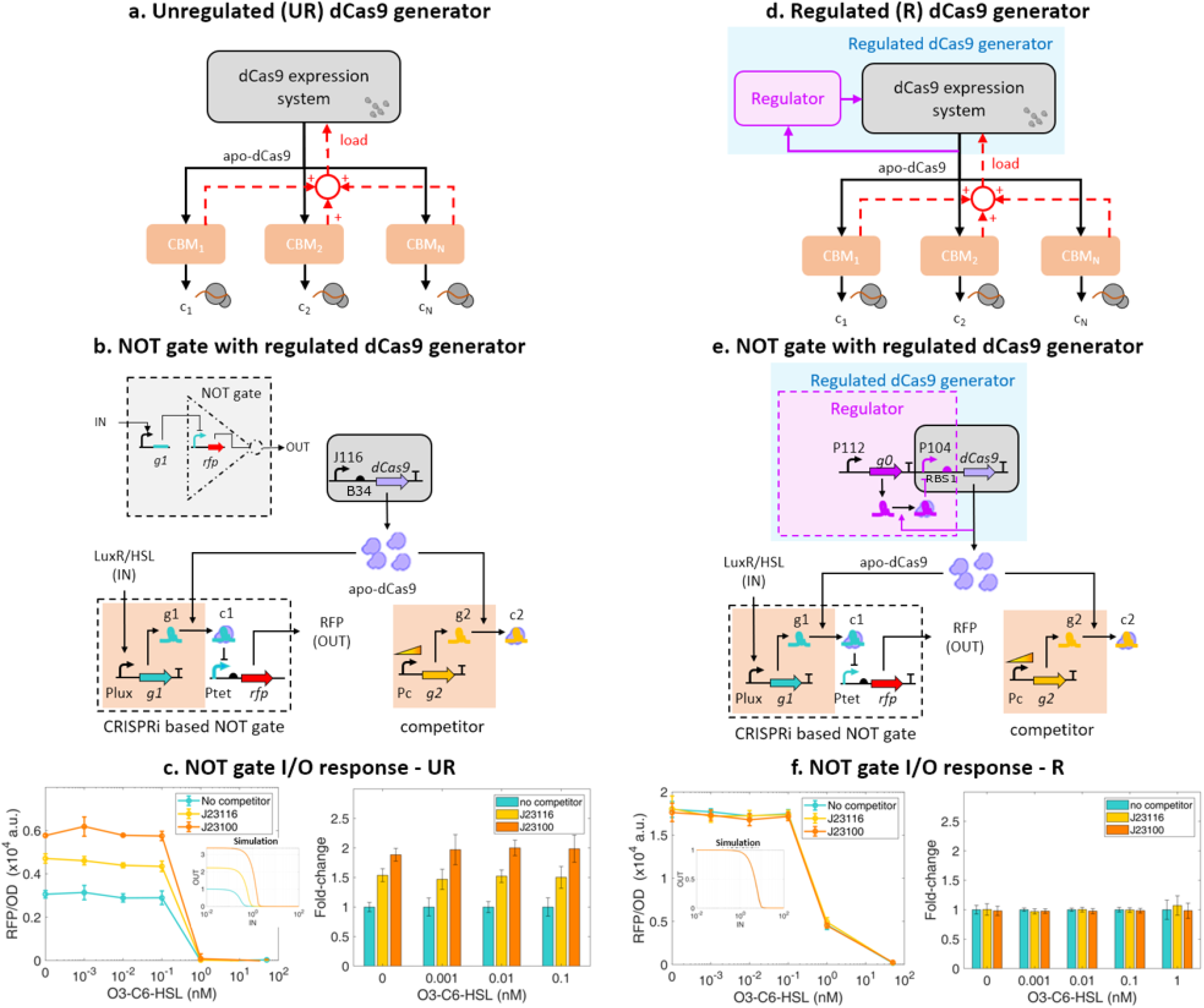
Regulated dCas9 generator mitigates effects of dCas9 loading on a CRISPRi-based NOT gate. (**a**) Unregulated dCas9 generator. A CRISPRi-based module (CBM) comprises a sequencespecific sgRNA, takes apo-dCas9 as an input, and produces the sgRNA-dCas9 complex “c” as a transcriptional repressor output. The CBM may have other regulatory inputs that control the expression of the sgRNA. The effect of competition for dCas9 exerted by different sgRNAs is represented as a load that each CBM applies to the dCas9 generator. (**b**) CRISPRi-based NOT gate with an unregulated dCas9 generator. The NOT gate is composed of a CBM comprising sgRNA g_1_, giving complex c_1_ as an output, and of a genetic module that takes c_1_ as input repressor and gives RFP protein as an output. Here, g_1_’s expression is regulated by HSL/LuxR. A second CBM expressing competitor sgRNA g_2_ with variable promoter Pc is either present or absent. The sgRNA expression cassettes were placed on a low copy number (~5) plasmid, dCas9 generators in a medium copy number (~20) plasmid, and the genetic module expressing RFP was borne on a high copy number (~200) plasmid. Details about parts and plasmids are reported in Supplementary Note 2. (**c**) Effect of expressing sgRNA competitor g_2_ on the NOT gate I/O response with unregulated (UR) dCas9 generator. Turquoise line represents I/O response in the absence of competitor, while yellow and organge lines represent system I/O responses in which the expression of competitor sgRNA is driven by a weaker (BBa_J23116) or stronger promoter (BBa_J23100), respectively (See Supplementary Note 2, Supplementary Table 6). Inset shows simulations based on the ODE model of the system described in Supplementary Equations (30)-(32) with parameters in Supplementary Table 9. (**d**) Block diagram representation of the regulated (R) dCas9 generator. The generator comprises a dCas9 expression system whose operation is modulated by a regulator as a function of the apo-dCas9 level. (**e**) The regulator is implemented by a constitutively expressed sgRNA g_0_ that targets dCas9 promoter for repression. dCas9 promoter is P104 and the RBS is RBS1, both stronger than those of the unregulated generator where dCas9 promoter is BBa_J23116 (shown as J116) and the RBS is BBa_B0034 (shown as B34). Details about parts and plasmids are reported in Supplementary Note 2 and Supplementary Note 5. (**f**) Effect of competitors on NOT gate I/O response with regulated (R) dCas9 generator with varying competitor levels as in panel (c). Inset shows simulations of the ODEs listed in Supplementary Equations (33)-(35) with parameters in Supplementary Table 9. Error bars represent the standard deviation of at least 3 biological replicates. Fold change of repression levels normalized on the no competitor data are computed as described in Online Methods Equation 1.

We created an experimental model system that recapitulates the dCas9 loading problem (Figure 1b). This model system includes a CRISPRi-based logic NOT gate, which is a key building block of any CRISPRi-dCas9 circuit, constitutive expression of a competitor sgRNA with variable promoter strength, and a dCas9 generator. The CRISPRi-based NOT gate is constituted of a CRISPRi-based module (CBM) with a primary sgRNA (g_1_) expressed through a HSL-inducible promoter and a genetic module expressing red fluorescent protein (RFP) as an output. The sgRNA g_1_ represses RFP’s transcription through CRISPRi (Supplementary Note 1, Supplementary Table 1). A second CBM contains sgRNA g_2_ (competitor) expressed by a constitutive promoter with varying strength (Supplementary Note 2, Supplementary Table 6). This competitor sgRNA plays the role of any additional sgRNA that may become expressed in a system, such as from other logic gates [3, 12]. We designed the competitor sgRNA g_2_ to target a DNA sequence not present in the circuit nor in the bacterial genome (as predicted by Benchling’s sgRNA desing tool, see Supplementary Note 1), in order to avoid further interactions that could confound analysis, such as metabolic changes due to binding the bacterial chromosome [13]. Both the NOT gate and competitor sgRNA are expressed from low copy number plasmid pSB4C5 with a pSC101 ORI (~5 copies, Supplementary Note 2). The dCas9 generator expresses dCas9 from a constitutive expression system (described in Supplementary Note 2, Supplementary Figure 1, also see [14]), to produce dCas9 at a level that is sufficient to enable complete repression of individual target promoters by sgRNAs without substantially affecting growth rate (Supplementary Note 3). We call this dCas9 expression system the unregulated (UR) dCas9 generator. With this generator, when the competitor sgRNA is expressed, the input/output (I/O) response of the CRISPRi-based NOT gate changes by up to two-fold, depending on the input level (Figure 1c). This is in accordance with simulations of an ordinary differential equation (ODE) model that captures the binding reactions between dCas9 and sgRNAs and between the dCas9-sgRNA complex and the corresponding target promoter (Figure 1c inset and Supplementary Equations (30)-(32)). Although these loading effects may, in principle, be mitigated by increased levels of dCas9, it is practically difficult to increase dCas9 level as this causes severe growth defects [6, 15, 16]. Less toxic mutations of dCas9 protein have been considered. Yet, even at the allowed higher dCas9 levels, the effects of dCas9 loading remain prominent [8].

Here, we introduce a regulated dCas9 generator that maintains the intended I/O response of CRISPRi-based logic gates independent of loads by competitor sgRNAs, while keeping dCas9 at sufficiently low levels to avoid growth defects (Figure 1d). This generator comprises regulation of the apo-dCas9 pool, which is achieved by an sgRNA g_0_ that represses dCas9’s promoter by CRISPRi (Figure 1e). In this design, when a competitor sgRNA g_2_ is expressed, it loads dCas9 such that the level of apo-dCas9 drops. This drop, in turn, reduces the level of dCas9-g_0_ complex, thereby both releasing dCas9 and de-repressing the dCas9 promoter. As a result, the level of apo-dCas9 increases, and this increase can balance the initial drop due to loading by the competitor sgRNA. When the dCas9 promoter is fixed, our mathematical model shows that the sensitivity of apo-dCas9 level to the expression rate of the competitor sgRNA g_2_ can be arbitrarily diminished by picking a sufficiently large expression rate for sgRNA g_0_ (Supplementary Note 4.3). This sensitivity reduction, in turn, theoretically mitigates the effects of competitor sgRNA g_2_’s expression on the NOT gate I/O response (simulations in Supplementary Figure 6). This is confirmed by the experimental data in Supplementary Figure 7b (left side and middle panels). These data also show that increased g_0_’s production rate can reduce the extent to which sgRNA g_1_ represses its target, due to lower apo-dCas9 concentration. The mathematical model indicates that the extent of repression of each sgRNA can be improved without sacrificing robustness by increasing dCas9 protein production rate when g_0_’s expression rate is sufficiently large (simulations in Supplementary Figure 6). This is validated by the experimental results in Supplementary Figure 7b (middle and right-side panels). Therefore, in the regulated dCas9 generator we also increased both dCas9’s promoter and RBS strengths compared to those of the unregulated generator (compare Figure 1b (UR) with Figure 1e (R)). When using this regulated dCas9 generator, the CRISPRi-based NOT gate maintains the same I/O response independent of the competitor sgRNA (Figure 1f). Specifically, while with an unregulated dCas9 generator the fold change of the output of the NOT gate is up to two-fold (Figure 1c), the fold change is inappreciable when a regulated dCas9 generator is used (Figure 1f). This is in accordance with simulations of an ODE model that includes regulation of dCas9 concentration (Figure 1f inset and Supplementary Equations (33)-(35)).

The load mitigation property of the regulated dCas9 generator generalizes to when sgRNAs are expressed from plasmids with high copy numbers, to alternative commonly used *E. coli* strains, and to different input regulators for the NOT gate (See Supplementary Figure 9 and Supplementary Figure 10). Specifically, we implemented the system of Figure 1b by placing both the CRISPRi-based NOT gate and competitor sgRNA g_2_ on a high copy number plasmid (pSC101E93G, ~ 84 copies, see Supplementary Note 6). This renders substantially larger dCas9 loads than those encountered in Figure 1, in which both the NOT gate and the competitor sgRNA are on a low copy number plasmid (~ 5 copies). These experiments were also carried in TOP10 strain as opposed to NEB10B strain. In these conditions, when the competitor sgRNA g_2_ is expressed and the unregulated dCas9 generator is used, the I/O response of the NOT gate changes by up to 25-fold, a much larger change than that observed in Figure 1. Nevertheless, when the regulated dCas9 generator is used, the change in the NOT gate I/O response is barely appreciable (~ 1.2 fold) in the same competitor conditions (Supplementary Figure 9). We then changed the input regulator to the NOT gate and, instead of using LuxR/HSL, we employed TetR/aTc. With this regulator, when a competitor sgRNA results in up to 13-fold change in the NOT gate’s I/O response with unregulated dCas9 generator, it gives an inappreciable change when the regulated generator is used (Supplementary Figure 10).

To demonstrate that the ability of the regulated dCas9 generator to mitigate the effects of dCas9 loading is not specific to a single CRISPRi-based NOT gate, we built a layered logic circuit constituted of two NOT gates arranged in a cascade (Figure 2a). The NOT gate cascade is a prototypical example of layered gates and is ubiquitous in circuits computing sophisticated logic functions [1, 3, 6, 12, 17, 18]. We thus constructed the cascade shown in Figure 2b, in which the LuxR/HSL input activates expression of an sgRNA g_1_, which, in turn, represses the expression of sgRNA g_3_ through CRISPRi. This sgRNA then represses, through CRISPRi, a constitutive promoter expressing the RFP output protein. As before, a competitor sgRNA g_2_ was included or omitted from the system. Also, while in the CRISPRi-based NOT gate (Figure 1b) all sgRNAs are expressed from a low copy number plasmid, we placed the cascade sgRNAs, RFP, and competitor sgRNA on higher copy number plasmid pSC101E93G (~84 copies) (Supplementary Note 6, Supplementary Table 4). This allows us to assess the regulated dCas9 generator performance in a situation of higher dCas9 loads. The I/O response of the cascade in Figure 2b was measured with or without the competitor sgRNA and with either the unregulated or regulated dCas9 generators. The I/O response of the cascade shows approximately a 4-fold change for low induction levels when the competitor sgRNA is added and the unregulated dCas9 generator is used (Figure 2c). By contrast, the cascade’s I/O response shows no appreciable change upon addition of the competitor when the regulated dCas9 generator is used, in accordance with the ODE model simulations (Figure 2d).

**Figure 2:**
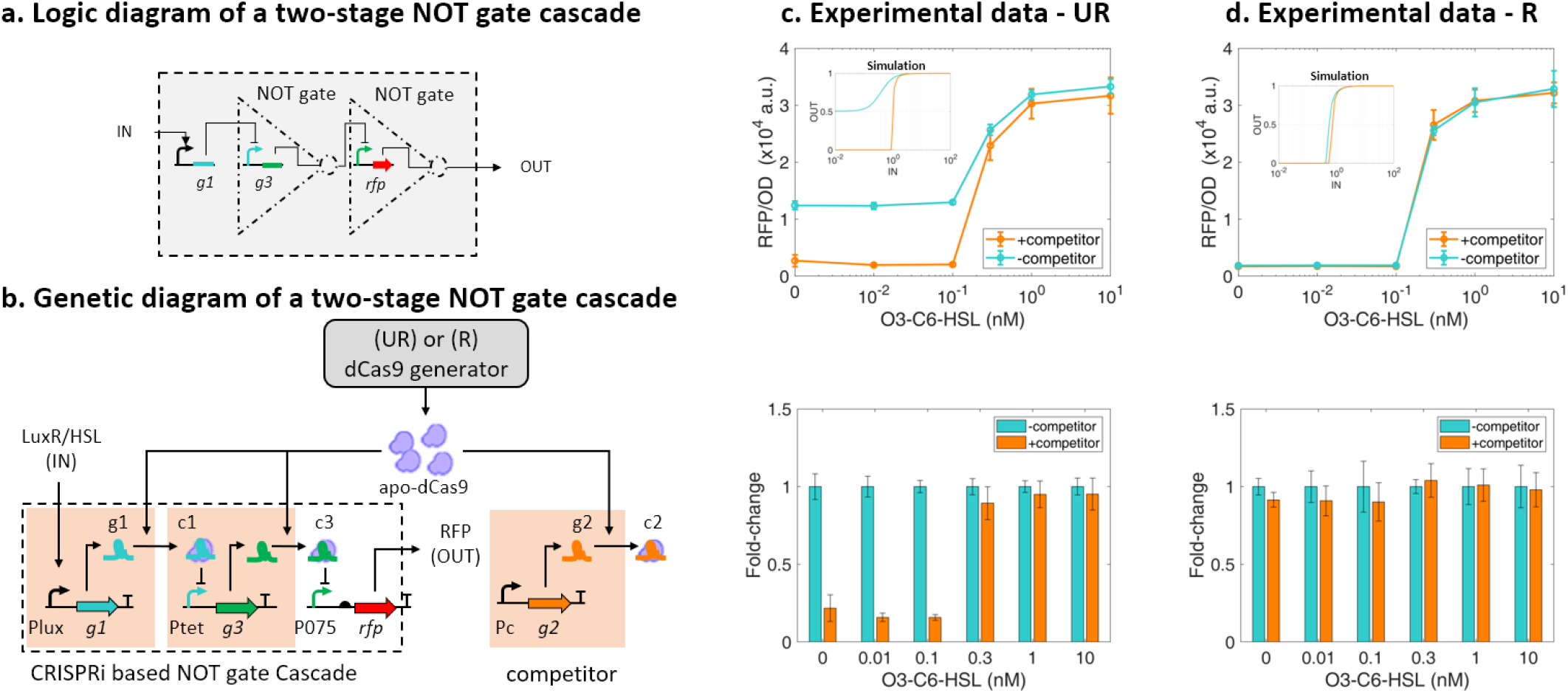
Regulated dCas9 generator mitigates effects of dCas9 loading on layered CRISPRi-based circuits. (**a**) Logic diagram of a NOT gates two-stage cascade based on CRISPRi modules. (**b**) Genetic circuit implementation used for both unregulated (UR) and regulated (R) versions of the dCas9 generator. The first NOT gate is constituted of a CBM comprising sgRNA g_1_, taking LuxR/HSL as input and giving repressive complex c_i_ as an output. The second NOT gate is constituted of a CBM comprising sgRNA g_3_ and of a genetic module expressing RFP from a promoter repressed by complex c_3_. The unregulated (UR) dCas9 generator is as in Figure 1b while the regulated (R) dCas9 generator is as in Figure 1e. Details about parts and plasmids are reported in Supplementary Note 2. (**c**) Effect of competitor sgRNA g_2_ on the cascade’s I/O response with unregulated (UR) dCas9 generator. Turquoise line represents system I/O response in the absence of the competitor, while organge line represents system I/O response in which the expression of competitor sgRNA is driven by a strong promoter (P108), (See Supplementary Note 2, Supplementary Table 6). Inset shows simulations based on ODEs listed in Supplementary Equations (36)-(38) with parameters in Supplementary Table 9. (**d**) Effect of competitor sgRNA g_2_ on the cascade’s I/O response with regulated (R) dCas9 generator. Inset shows simulations with the ODEs in Supplementary Equations (39)-(41) with parameters in Supplementary Table 9. Error bars in the plots represent the standard deviation of at least 3 biological replicates. Fold change of repression levels normalized to the no competitor data are reported as described in Online Methods Equation 1.

Taken together, our data demonstrate that the regulated dCas9 generator effectively removes interference among otherwise orthogonal sgRNAs, which results from sharing a limited pool of dCas9. Thus, the regulated dCas9 generator restores true orthogonality among sgRNA-promoter pairs to support creation of increasingly complex transcriptional programs. The regulated dCas9 generator is implemented in its own dedicated plasmid (Supplementary Note 2, Supplementary Figure 1) and, as such, can be easily transported across compatible bacterial strains and applications of CRISPRi-dCas9 systems. Feedback regulation systems have been designed to enhance robustness of bacterial genetic circuits to loading of shared gene expression resources, i.e., the ribosome [19–21]. However, none of these approaches is directly applicable to regulate apo-dCas9 level as they either use ribosome-specific parts [21], require the protein to be regulated (dCas9 in our case) to act as a transcriptional activator [19], or to be able to sequester a transcriptional activator [20]. By contrast, the dCas9 regulator is simple, compact, and exploits directly the ability of dCas9 to function as a transcriptional repressor. When expressing multiple sgRNAs from the chromosome, i.e., in one copy, reduced loading effects are expected [22] and a regulated dCas9 generator may not be required in such cases. However, we have shown that for CRISPRi-dCas9 systems constructed on plasmids, loading effects are prominent even at low plasmid copy number (~5 copies, Figure 1c). In these cases, it is expected that a regulated dCas9 generator will be required in order to ensure that multiple sgRNAs can concurrently and independently control their targets.

## ONLINE METHODS

Materials and methods are reported in the online version of the paper.

### Strain and growth medium

Bacterial strain *E. coli* NEB10B (NEB, C3019I) was used in genetic circuit construction and characterization. The growth medium used in construction was LB broth Lennox. The growth medium used in characterization was M9 medium supplemented with 0.4 % glucose, 0.2 % cascamino acids, and 1 mM thiamine hydrochloride. Appropriate antibiotics were added according to the selection marker of a genetic circuit. Final concentration of ampicillin, kanamycin and chloramphenicol are 100, 25, and 12.5 μgmL^−1^, respectively.

### Genetic circuit construction

The genetic circuit construction was based on Gibson assembly method [23]. DNA fragments to be assembled were amplified by PCR using Phusion High-Fidelity PCR Master Mix with GC Buffer (NEB, M0532S), purified with gel electrophoresis and Zymoclean Gel DNA Recovery Kit (Zymo Research, D4002), quantified with the nanophotometer (Implen, P330), and assembled with Gibson assembly protocol using NEBuilder HiFi DNA Assembly Master Mix (NEB, E2621S). Assembled DNA was transformed into competent cells prepared by the CCMB80 buffer (TekNova, C3132). Plasmid DNA was prepared by the plasmid miniprep-classic kit (Zymo Research, D4015). DNA sequencing used Quintarabio DNA basic sequencing service. The list of primers and constructs is in Supplementary Table 5.

### Microplate photometer measurements

Overnight culture was prepared by inoculating a −80°C glycerol stock in 800 μL growth medium per well in a 24-well plate (Falcon, 351147) and grew at 30 °C, 220 rpm in a horizontal orbiting shaker for 13 h. Overnight culture was first diluted to an initial optical density at 600 nm (OD_600nm_) of 0.001 in 200 μL growth medium per well in a 96-well plate (Falcon, 351172) and grew for 2h to ensure exponential growth before induction. The 96-well plate was incubated at 30 °C in a Synergy MX (Biotek, Winooski, VT) microplate reader in static condition and was shaken at a fast speed for 3s right before OD and fluorescence measurements. Sampling interval was 5 min. Excitation and emission wavelengths to monitor RFP fluorescence were 584 and 619 nm, respectively. To ensure exponential growth, cell culture was diluted with fresh growth medium to OD_600nm_ of 0.01 when OD_600nm_ approaches 0.12 at the end of one batch. Multiple batches were conducted until gene expression reaches steady state. Growth rates were computed from the last batch of each experiment.

### Quantification of competition effects

To quantify competition effects, fold-change of a system at a given induced condition j to dCas9 competition was calculated by taking the ratio of the RFP/OD value of the system with competitor sgRNA (g_2_) to the corresponding system bearing no competitor sgRNA (e.g. pOP69 and pCL87, Supplementary Figure 9c):

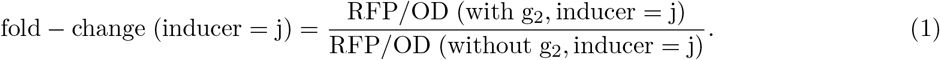

## Acknowledgements

The authors want to thank Pin-Yi Chen (Massachusetts Institute of Technology, USA) for the modeling insights throughout the initial phase of the study and Michela Casanova (University of Pavia, Italy) for the assistance with preliminary experiments and cloning activities.

## Author contributions

H.H., M.B., Y.Q., and P.C. designed constructs and performed experiments. Y.Q. and M.B. performed simulations and mathematical analyses. D.D.V., M.B., Y.Q., and H.H. wrote the paper and analyzed the data. P.M. and L.P. supervised the activities of M.B.; D.D.V. designed the research.

## Supplementary Information

### Supplementary Note 1 DNA sequences

DNA sequences of the plasmids listed in Supplementary Table 1 can be found in Supplementary Data. The guide sequences of all sgRNAs are listed in Supplementary Table 2. All designs of sgRNA sequences were aided with the CRISPR Guide RNA Design Tool of Benchling (Benchling.com).

**Supplementary Table 1:**
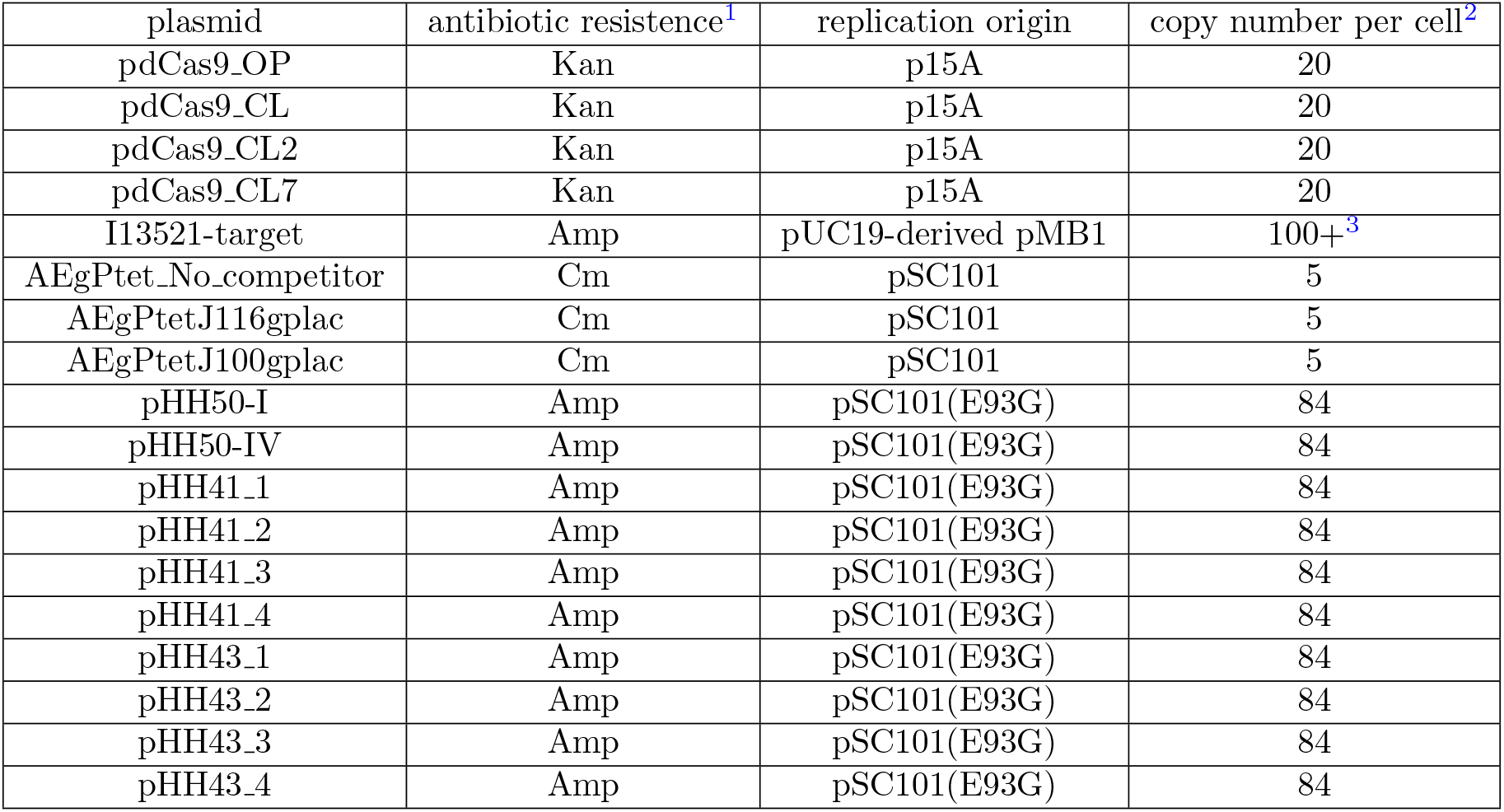
List of plasmids used in this work.

| plasmid | antibiotic resistance<sup>1</sup> | replication origin | copy number per cell<sup>2</sup> |
| --- | --- | --- | --- |
| pdCas9_OP | Kan | p15A | 20 |
| pdCas9_CL | Kan | p15A | 20 |
| pdCas9_CL2 | Kan | p15A | 20 |
| pdCas9_CL7 | Kan | p15A | 20 |
| I13521-target | Amp | pUC19-derived pMB1 | 100+<sup>3</sup> |
| AEgPtet_No_competitor | Cm | pSC101 | 5 |
| AEgPtetJ116gplac | Cm | pSC101 | 5 |
| AEgPtetJ100gplac | Cm | pSC101 | 5 |
| pHH50-I | Amp | pSC101(E93G) | 84 |
| pHH50-IV | Amp | pSC101(E93G) | 84 |
| pHH41.1 | Amp | pSC101(E93G) | 84 |
| pHH41.2 | Amp | pSC101(E93G) | 84 |
| pHH41.3 | Amp | pSC101(E93G) | 84 |
| pHH41.4 | Amp | pSC101(E93G) | 84 |
| pHH43.1 | Amp | pSC101(E93G) | 84 |
| pHH43.2 | Amp | pSC101(E93G) | 84 |
| pHH43.3 | Amp | pSC101(E93G) | 84 |
| pHH43.4 | Amp | pSC101(E93G) | 84 |

**Supplementary Table 2:**
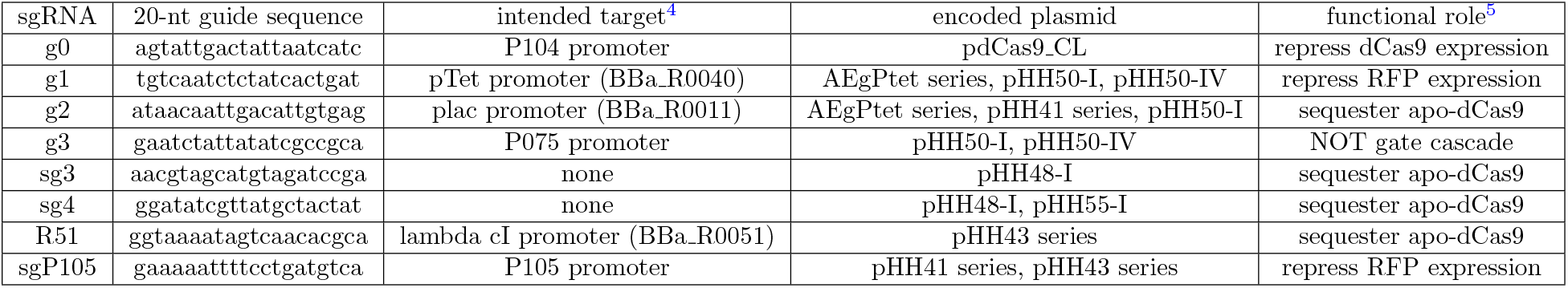
List of all sgRNAs used in this work.

### Supplementary Note 2 Genetic constructs in the main text

A genetic construct (Figure 1b), in which the CRISPRi-based NOT gate competes with a different amount of the competitor sgRNA g2 for apo-dCas9 proteins produced by the unregulated dCas9 generator, was implemented with the constructs pOP69, pOP70, and pOP71 listed in Supplementary Table 3. Specifically, each construct is composed of the indicated three plasmids. The pdCas9_OP plasmid encodes the unregulated dCas9 generator and is the plasmid j116-dcas9-3k3 (BBa_J107202) from [S1]. The NOT gate is implemented as a CRISPRi-based module (CBM) in an AEgPtet plasmid and its target cassette is in the I13521-target plasmid. The competitor sgRNA g2 cassette is absent in the AEgPtet_No_competitor plasmid of the construct pOP69, but is present in the AEgPtet_116gPlac and AEgPtet_100gPlac plasmids of the constructs pOP70 and pOP71, respectively. The AEgPtet_116gPlac and AEgPtet_100gPlac plasmids use BBa_J23116 and BBa_J23100 promoters, respectively, to transcribe the competitor sgRNA g2. Similarly, the genetic circuit in Figure 1e was implemented with the constructs pCL87, pCL88, and pCL89, respectively. The pdCas9_CL plasmid encodes the regulated dCas9 generator and is constructed by Gibson assembly as shown in the Supplementary Table 5.

**Supplementary Table 3:**
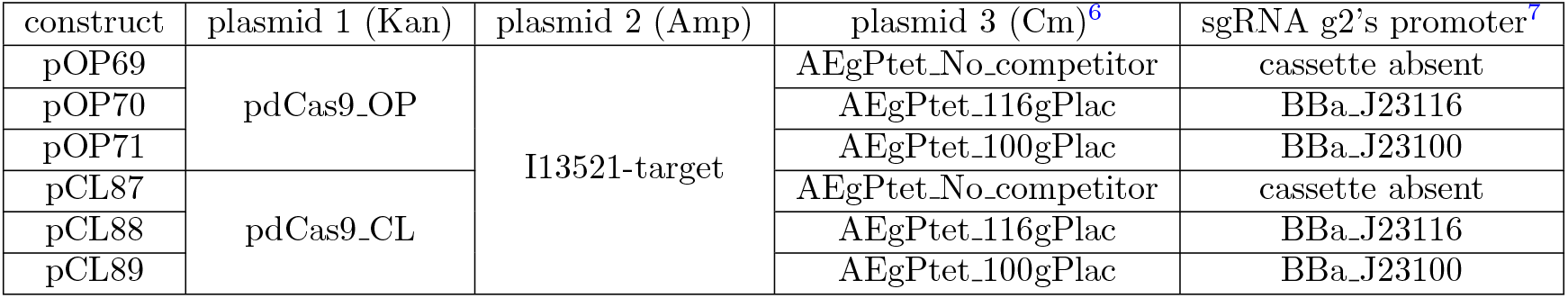
List of the constructs used in Figure 1. Each construct was obtained by cotransformation of the three indicated plasmids into *E. coli* NEB10B strain. The component plasmids 1, 2, and 3 have antibiotic resistance kanamycin, ampicillin, and chloramphenicol, respectively, and have the origin of replication as p15A [S2], pUC19-derived pMB1 [S3], and pSC101 [S2], respectively.

The genetic cirucit (Figure 2b), in which the CRISPRi-based NOT gate cascade operates with and without the competitor sgRNA g2 for apo-dCas9 proteins produced by the unregulated dCas9 generator, was built as the constructs pOP64 and pOP65, respectively, as listed in Supplementary Table 4. Specifically, each construct is composed of the two indicated plasmids. pHH50-IV encodes the NOT gate cascade and does not encode the competitor sgRNA. pHH50-I is derived from pHH50-IV by introducing the competitor sgRNA g2 in downstream of the P108 promoter of pHH50-IV. The P108 promoter is adopted from the Ec-TTL-P108 promoter [S4]. Similarly, the genetic circuits using the regulated dCas9 generator with or wiothout the competitor sgRNA were built as the constructs pCL82 and pCL83, respectively. Gibson assembly and maps of the pHH50-I and pHH50-IV plasmids are shown in the Supplementary Table 5 and in the Supplementary Figure 3, respectively.

**Supplementary Table 4:**
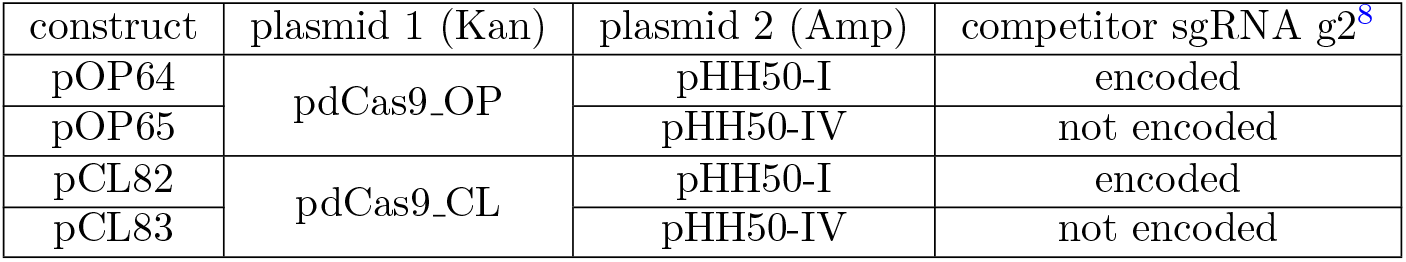
List of the constructs used in Figure 2. Each construct results from concurrent transformation of the indicated two plasmids into *E. coli* NEB10B strain. The plasmids 1 and 2 have antibiotic resistance kanamycin and ampicillin, respectively, and the origin of replication as p15A and pSC101(E93G) [S2], respectively.

| construct | plasmid 1 (Kan) | plasmid 2 (Amp) | competitor sgRNA g2<sup>8</sup> |
| --- | --- | --- | --- |
| pOP64 | pdCas9_OP | pHH50-I | encoded |
| pOP65 |  | pHH50-IV | not encoded |
| pCL82 | pdCas9_CL | pHH50-I | encoded |
| pCL83 |  | pHH50-IV | not encoded |

**Supplementary Table 5:** List of plasmids which were assembled by Gibson assembly and used in Supplementary Table 3 and Supplementary Table 4. Each fragment was prepared by the indicated forward and reverse primers and the DNA template.

| Plasmid | Fragment | Forward | Reverse | DNA Template | Size (bp) |
| --- | --- | --- | --- | --- | --- |
| pdCas9_CL | 1 | P264 | P312 | P112_sgRNA_P104_dCas9_B34 | 2143 |
|  | 2 | P311 | AH1prR | P112_sgRNA_P104_dCas9_B34 | 2186 |
|  | 3 | AH2prF | P263 | P112_sgRNA_P104_dCas9_B34 | 1318 |
|  | 4 | P262 | GFP-VR | P112_sgRNA_P104_dCas9_B34 | 1644 |
| pHH50-I | 1 | Load_F2 | P662 | pHH48-I #2 | 2424 |
|  | 2 | P663 | P664 | 2ndStage_RFP | 120 |
|  | 3 | P665 | P730 | 2ndStage_RFP | 534 |
|  | 4 | P731 | P667 | pHH48-I #2 | 1090 |
|  | 5 | P668 | P367 | AEgPtetYgpComp-double-knob | 1129 |
|  | 6 | P669 | Load_R2 | pHH48-I #2 | 1855 |
| pHH50-IV | 1 | Load_F2 | P662 | pHH48-I #2 | 2424 |
|  | 2 | P663 | P664 | 2ndStage_RFP | 120 |
|  | 3 | P665 | P730 | 2ndStage_RFP | 534 |
|  | 4 | P731 | P667 | pHH48-I #2 | 1090 |
|  | 5 | P668 | P367 | AEgPtetYgpComp-double-knob | 1129 |
|  | 6 | P669 | P613 | pHH48-I #2 | 351 |
|  | 7 | P614 | Load_R2 | pHH48-I #2 | 1442 |

**Supplementary Table 6:** Promoters used in the main text with relative strength, as evaluated in [S4] and [S1], normalized to BBa_J23116.

| Promoter | Use | Strength | Plasmid | Reference |
| --- | --- | --- | --- | --- |
| BBa_J23116 | dCas9 expression in unregulated systems<br>g2 weak expression in NOT gate systems | 1 | pdCas9_OP<br>AEgPtet_116gPlac | [S4] |
| BBa_J23100 | g2 strong expression in NOT gate systems | 20 | AEgPtet_100gPlac | [S4] |
| P104 | dCas9 expression in regulated systems | 130 | pdCas9_R | [S4] |
| P112 | g0 expression in regulated systems | 258 | pdCas9_R | [S4] |
| P108 | g2 expresison in Cascade systems | 173 | pHH50-IV | [S4] |
| Ptet | RFP expression in NOT gate system<br>g3 expression in Cascades system | 34 | I13521-target<br>pHH50-I,pHH50-IV | [S4] |
| P075 | RFP expression in Cascade systems | 48 | pHH50-I, pHH50-IV | [S4] |
| Plux<sup>9</sup> | g1 expression in all the reported systems | 97 | AEgPtet_No_competitor<br>AEgPtet_116gPlac<br>AEgPtet_100gPlac<br>pHH50-I, pHH50-IV | [S4, S1] |

**Supplementary Figure 1:**
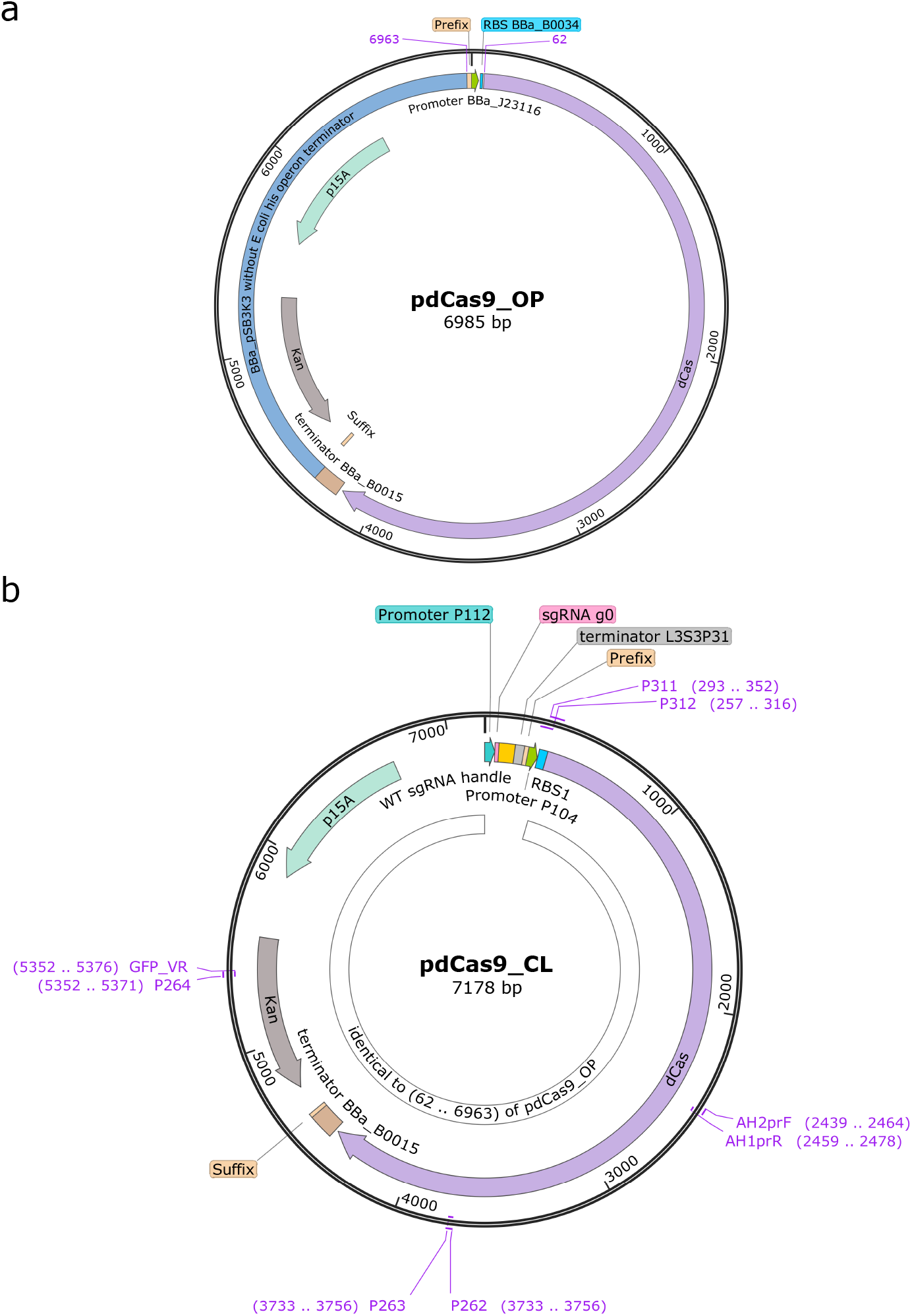
Maps of the pdCas9 OP and pdCas9 CL plasmids used in the constructs listed in Supplementary Table 3, Supplementary Table 4, and Supplementary Table 12. (a) The pdCas9 OP plasmid uses the promoter BBa_J23116 and the RBS BBa_B0034 to constitutively express dCas9 protein. The annotated map section from 62 to 6963 is identical to the pdCas9 CL plasmid. (b) The pdCas9 CL plasmid uses the regulator as shown in Figure 1e to control the expression of dCas9 protein. Specifically, the regulator comprises the strong promoter P104 and the strong RBS *RBS*1 to express dCas9 protein and uses the strong promoter P112 to constitutively transcribe the sgRNA g0, which targets the promoter P104 of the *dCas9* gene to render the regulation. The P104 and P112 promoters are adopted from the Ec-TTL-P104 and Ec-TTL-P112 promoters, respectively [S4]. The sgRNA g0 is composed of the 20-nt sgRNA g0 sequence, the wild-type (WT) sgRNA handle [S7], and the terminator L3S3P31 [S8]. Furthermore, the 20-nt sgRNA g0 sequence was designed with the CRISPR Guide RNA Design Tool of Benchling (Benchling.com). The cloning primers listed in Supplementary Table 5 are annotated as a purple text with the respective map position.

**Supplementary Figure 2:**
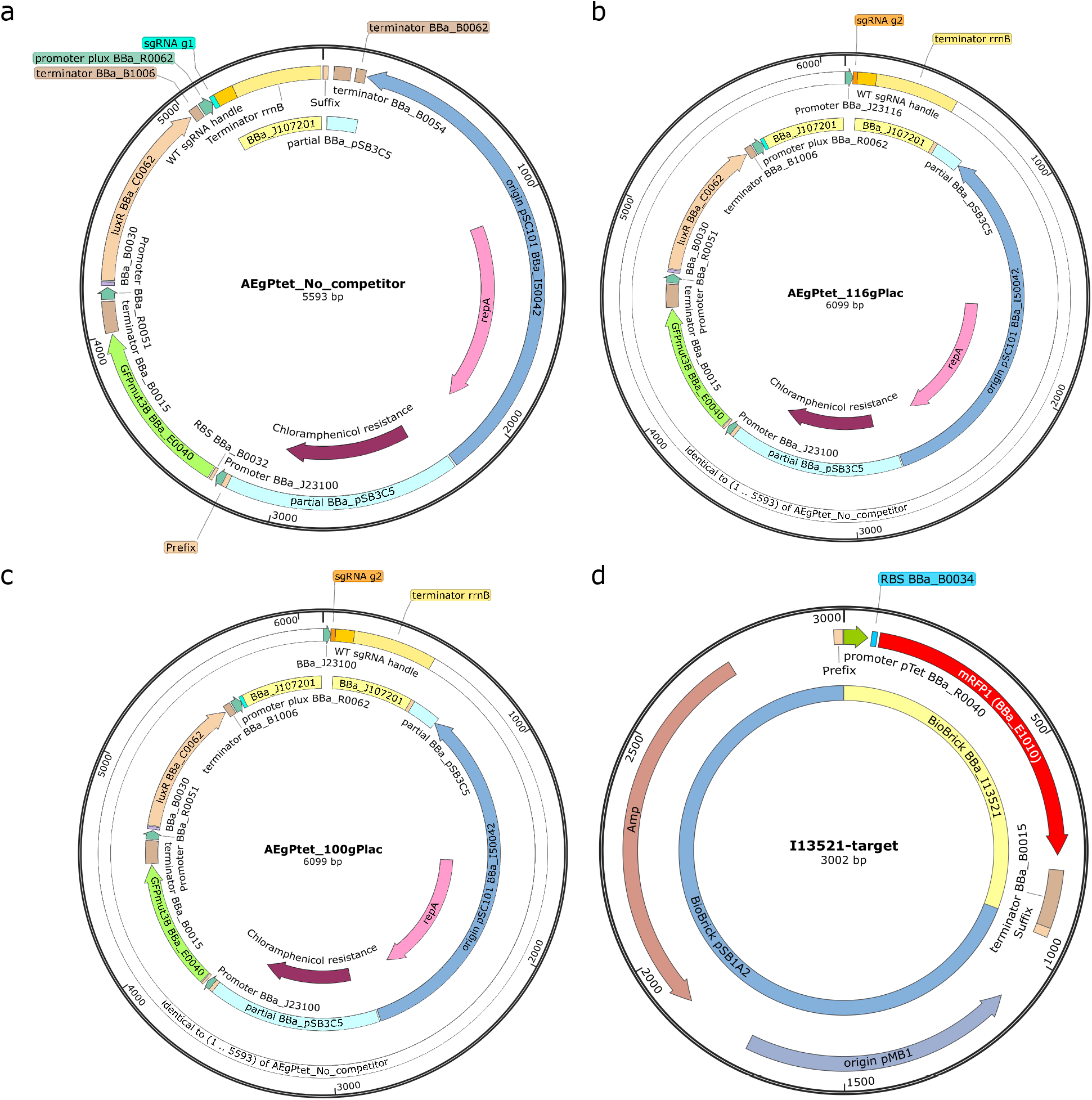
Maps of the AEgPtet and I13521-target plasmids which were used in the constructs listed in Supplementary Table 3. (a-c) In the AEgPtet plasmid, specifically, the NOT gate uses transcriptional activator LuxR and its effector HSL (i.e. LuxR/HSL) as the input and red fluorescent protein RFP as the output. LuxR/HSL activates the Plux promoter to transcribe the sgRNA g1 from the AEgPtet plasmid. The dCas9-sgRNA g1 complex represses the pTet promoter of the *mRFP1* gene in the I13521-target plasmid. AEgPtet_Nα.competitor does not encode the competitor sgRNA cassette. AEgPtet_116gPlac and AEgPtet_100gPlac plasmids use the BBa_J23116 and the BBa_J23100 promoters to constitutively transcribe the sgRNA g2, respectively. (d) The I13521-target plasmid uses the promoter pTet to constitutively express red fluorescence protein (mRFP1). The 20-nt guide sequences of the sgRNAs g1 and g2 were designed with the CRISPR Guide RNA Design Tool of Benchling (Benchling.com) to target the pTet and BBa_R0011 promoters, respectively, without predicted off-targets in the genome of *E. coli* K-12 strain. Note that the BBa_R0011 promoter is not used in any plasmid of this work. The sgRNA g1 and g2 have the common BBa_J107201 BioBrick part which includes the WT sgRNA handle and the terminator rrnB.

**Supplementary Figure 3:**
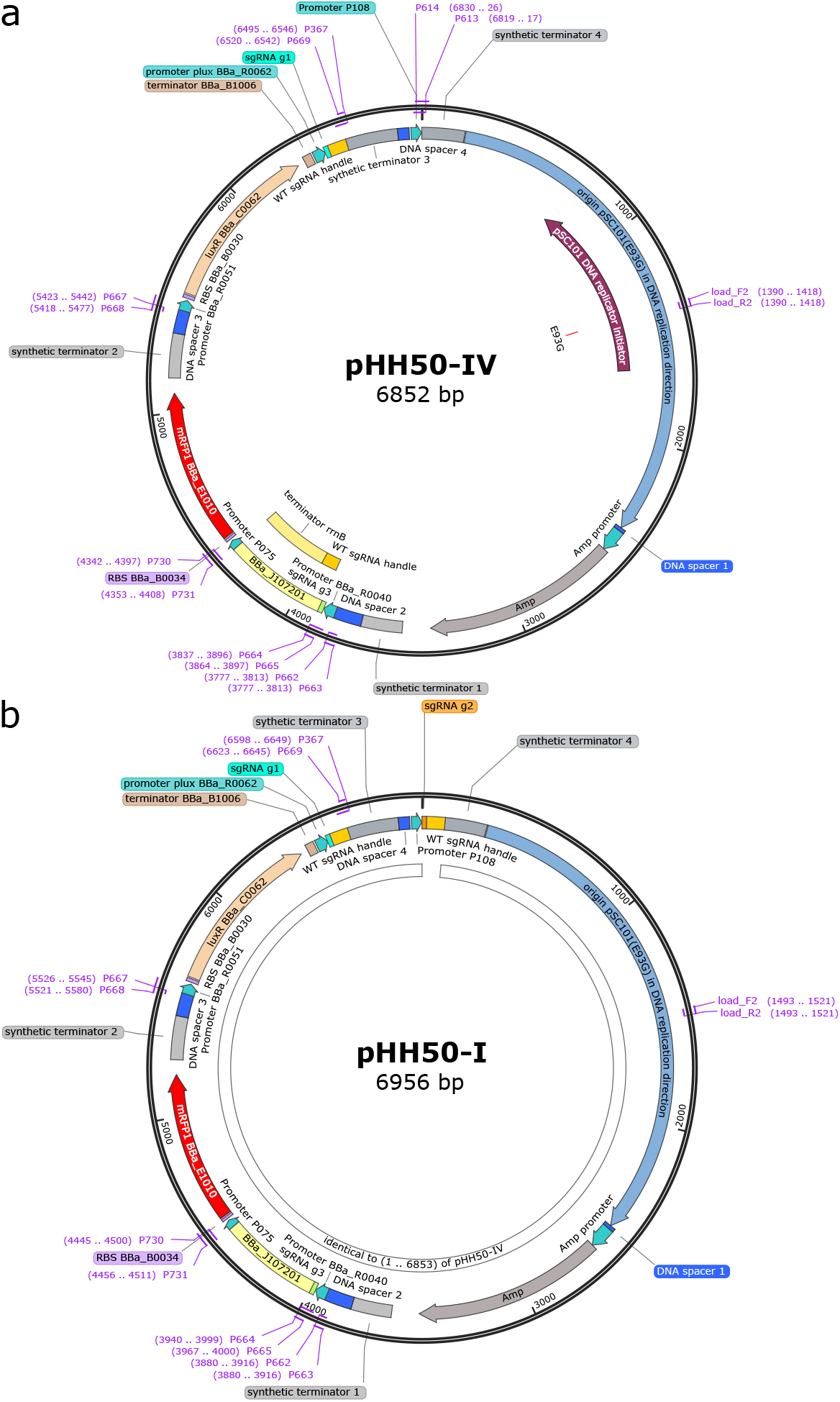
Maps of the pHH50-IV and pHH50-I plasmids which were used in the constructs listed in Supplementary Table 4. (a) The CRISPRi-based NOT gate cascade in Figure 2b is implemented in pHH50-IV and uses transcriptional activator LuxR and its effector HSL (i.e. LuxR/HSL) as the input and red fluorescent protein RFP as the output. Specifically, LuxR/HSL activates the plux promoter (BBa_R0062 on map position 6365) to transcribe the sgRNA g1 (on map position 6417) to target the pTet promoter. The first-stage NOT gate uses the pTet promoter (BBa_R0040 on map position 3816) to transcribe the sgRNA g3 (on map position 3871) to target the P075 promoter. The second-stage NOT gate uses the P075 promoter (on map position 4342) to express *mRFP1* gene as the output of the cascade. The P108 promoter is located immediately upstream of the synthetic terminator 4. No competitor sgRNA is encoded in this plasmid. The P075 and P108 promoters are adopted from the Ec-TTL-P075 and Ec-TTL-P108 promoters [S4]. (b) The pHH50-I plasmid encodes the 20-nt guide sequence of the competitor sgRNA g2 (on map position 1) and the WT sgRNA handle (on map position 21) in downstream of the P108 promoter (on map position 6910). The sgRNA g3 was designed in the same way as the sgRNAs g1 and g2 but only differs in the 20-nt guide sequence. The cloning primers listed in Supplementary Table 5 are annotated with purple text with the respective map position.

### Supplementary Note 3 Specific growth rates of the experiments in the main text

**Supplementary Figure 4:**
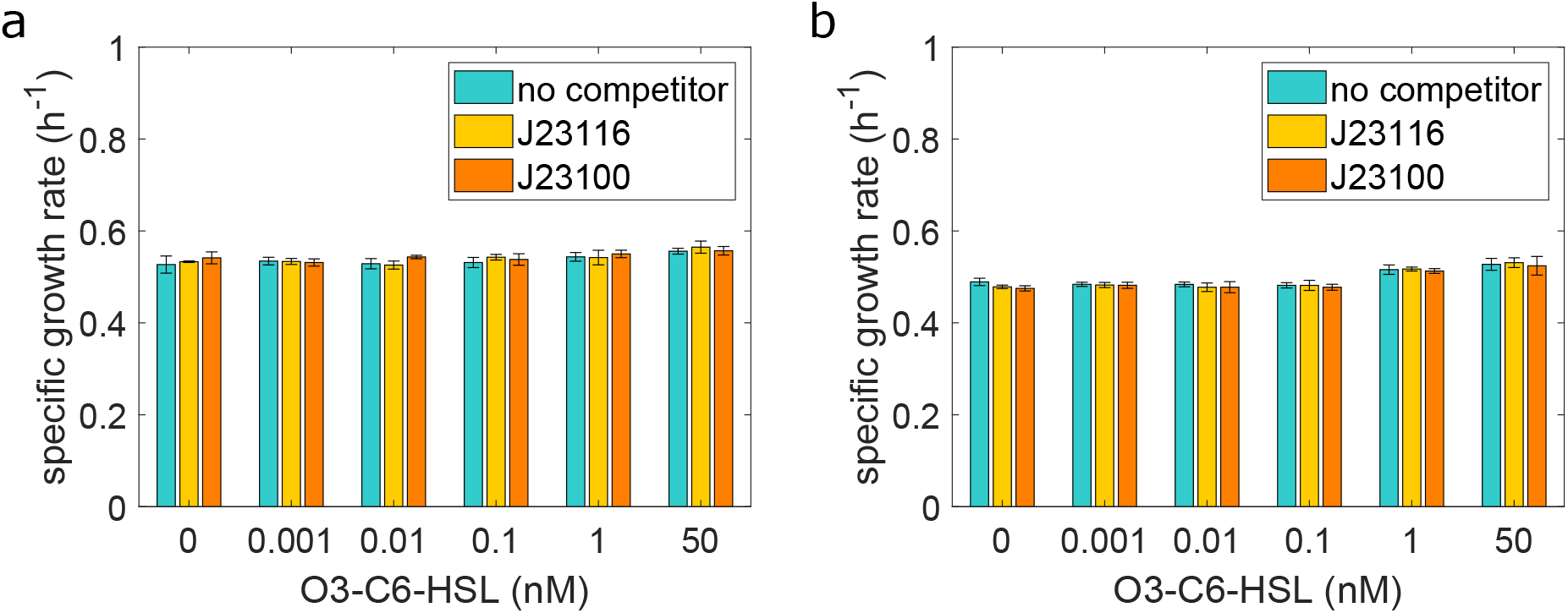
Specific growth rates observed in the experiments of the NOT gate circuit in Figure 1. (a) The rates at steady state were observed in the data of Figure 1c. (b) The rates at steady state were observed in the data of Figure 1f.

**Supplementary Figure 5:**
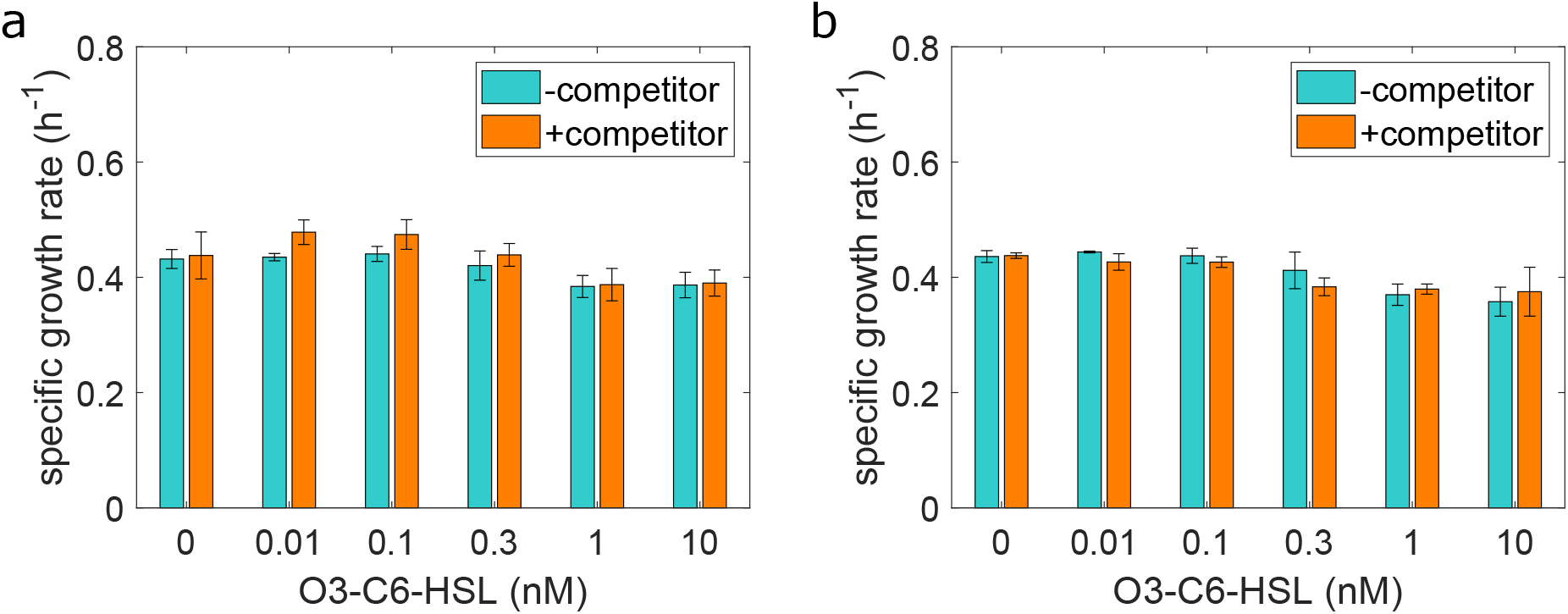
Specific growth rate observed in the experiments of the two-stage NOT gate cascade in Figure 2g. (a) The rates at steady state were observed in the data of Figure 2c. (b) The rates at steady state were observed in the data of Figure 2d.

### Supplementary Note 4 Modeling framework

In Supplementary Note 4.1, we first establish a general modeling framework for CRISPRi-based genetic circuits. The specific models for the NOT gate and the cascade are described in Supplementary Note 4.2. In Supplementary Note 4.3, we provide mathematical analysis to guide the regulated dCas9 generator design and parameter tuning. These analysis results are used to educate experimental choices.

#### Supplementary Note 4.1 A general modeling framework for CRISPRi-based genetic circuits

In this section, we first describe the dynamics of the sgRNAs and the complexes they form, and then establish models for the dynamics of dCas9 protein and other proteins in the circuit.

##### sgRNA dynamics

We consider a CRISPRi-based genetic circuit composed of a set of sgRNAs *g_i_*, where *i* takes value in an index set 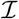. Each sgRNA *g_i_* can bind with apo-dCas9 (D) to form a dCas9-sgRNA complex *c_i_*. These sgRNAs further fall into two complementary subsets 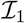 and 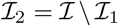. For sgRNA *g_i_* such that 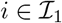, the complex *c_i_* can bind with its targeting site on promoter *p_t_i__* to form a complex *c_ii_*. Alternatively, for sgRNA *g_i_* such that 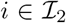, the complex *c_i_* does not have a targeting site. These biomolecular processes can be described by the following chemical reactions:

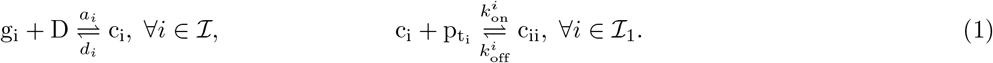

dCas9 protein can dissociate from sgRNA even when the dCas9-sgRNA complex is bound to DNA. To model this phenomenon, for 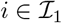, we also consider the reaction

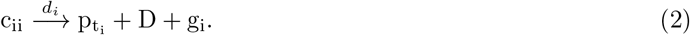

Let *p_i_* represent the promoter transcribing sgRNA *g_i_*, the production and decay of sgRNAs are described as:

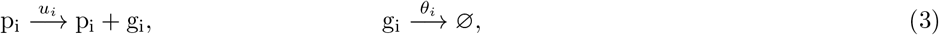

where *u_i_* is the synthesis rate constant of sgRNA from a single promoter and *θ_i_* is the degradation rate constant of sgRNA *g_i_*. The magnitude of the synthesis rate constant u_i_ increases with the strength of the promoter. Additionally, we take into account the fact that all species are diluted at rate constant *δ* due to cell growth:

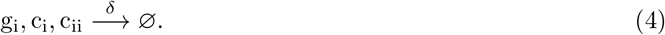

By mass action kinetics [S16], the chemical reactions in (1)-(4) can be modeled by the following ODEs:

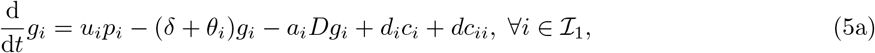

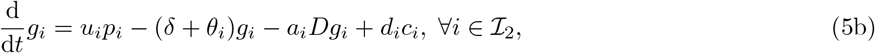

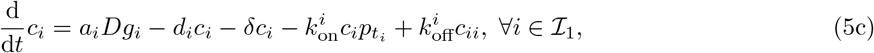

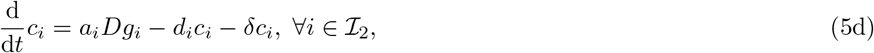

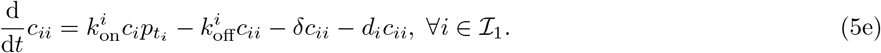

For 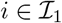, let 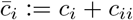 be the total amount of dCas9-sgRNA complex, using (5a) and (5c), we have

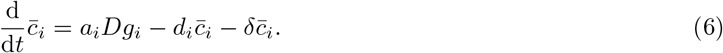

To obtain a reduced order model of (5) that facilitates analysis and educates design, we assume that binding and unbinding dynamics of the complexes *c_i_* and *c_ii_* are sufficiently fast compared to RNA and protein dynamics, and hence we assume their concentrations reach quasi-steady state (QSS). By setting the temporal derivatives in (5c)-(5e) to zero we obtain the QSS complex concentrations:

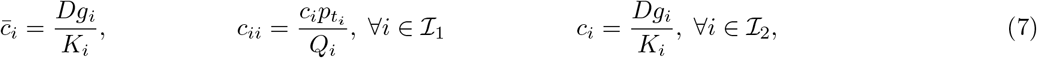

where

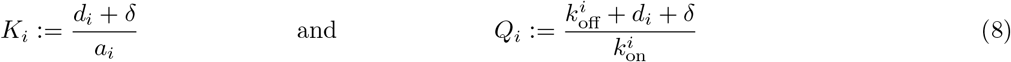

are the dissociation constants describing the binding between sgRNAs and apo-dCas9 protein, and between dCas9-sgRNA complex and the targeting promoter, respectively. Substituting (7) into (5a) and (5b), the dynamics of g_*i*_ can be re-written as:

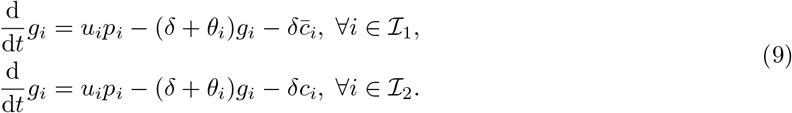

The pool of total dCas9 protein is shared by all sgRNAs in the circuit. Let *D_T_* represent the total dCas9 concentration, then dCas9 concentration follow the conservation law:

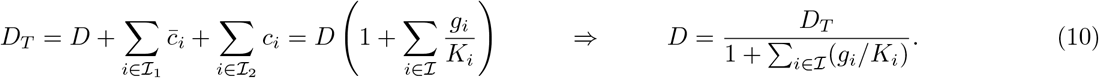

Substituting (10) into (7), the QSS concentrations of the complexes in (7) can be re-written as:

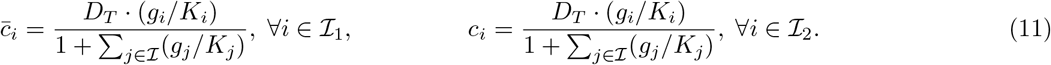

For 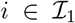, to find the extent of repression of *g_i_* on its target promoter *p_t_i__*, we need to compute the concentration of *c_ii_*. To this end, suppose that *t_i_* = *j* for some 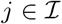, we note that the concentration of *p_j_* promoter follows the conservation law:

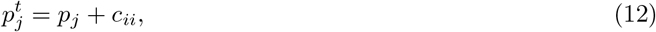

where 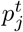 is the total concentration of the promoter driving the transcription of *g_j_*. Substituting the QSS of *c_ii_* in (7) into (12), we have

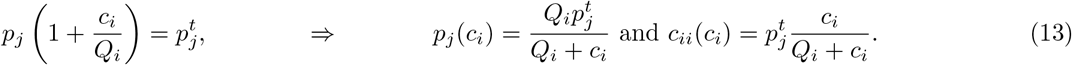

Using the QSS of *c_ii_* = *c_ii_*(*c_i_*) computed in (13), the QSS concentration of *c_i_* can be found through the equality

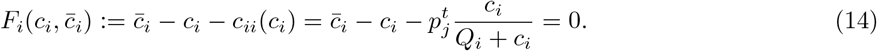

For a fixed and bounded 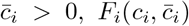 is a monotonically decreasing function of *c_i_* and it satisfies 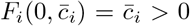 and 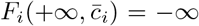. Thus, the equation 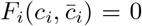 has a unique, positive solution 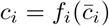. In particular, this solution can be computed as:

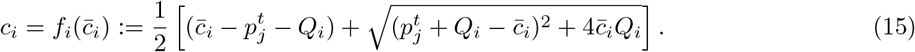

Substituting (15) into (13), we obtain the QSS concentration of *p_j_* available for transcription:

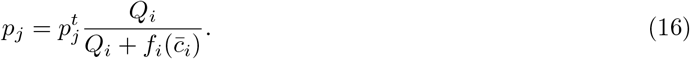

Substituting (16) and (11) into (9) and let *g* be the vector representing the concentrations of all sgRNAs, the dynamics of g_*i*_ can be written as:

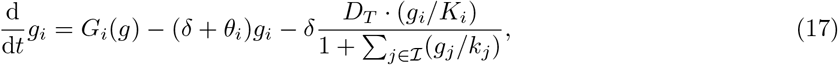

where

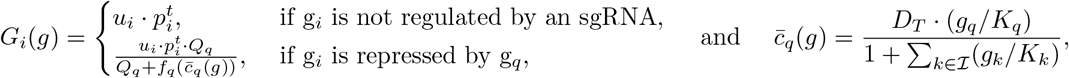

and function *f_q_*(·) is defined as in (15).

##### dCas9 protein dynamics

The synthesis and decay of dCas9 protein (D) can be modeled by the chemical reactions:

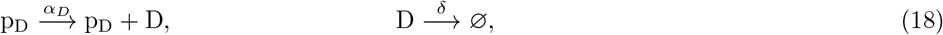

where *α_D_* is the synthesis rate constant of D from each copy of free promoter p_*D*_ driving dCas9 expression and *δ* is the dilution rate constant. The lumped parameter *α_D_* increases with, for example, the dCas9’s (i) promoter strength, (ii) plasmid copy number, and (iii) RBS strength. Based on mass action kinetics of the chemical reactions in (1), (2), and (18), we have

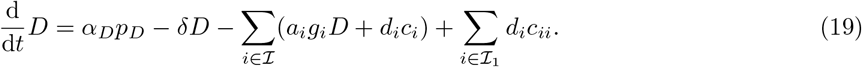

Substituting the QSS concentrations of the complexes in (7) into (19), the free dCas9 dynamics can be written as:

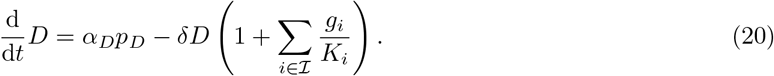

The free promoter concentration *p_D_* depends on whether dCas9 expression is regulated or not. Specifically, when dCas9 expression is unregulated, all promoters are available for transcription, and we set 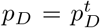. When dCas9 expression is repressed by sgRNA g_0_, the promoter concentration satisfies the conservation law 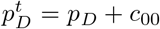. Similar to (16), using the QSS concentration of *c*_00_ derived in (7) and the relationship between 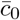 and *c*_0_ derived in (15), we have

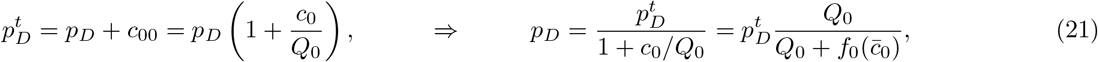

where 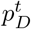 is the total concentration of promoter driving dCas9 expression and *Q*_0_ is dissociation constant between complex c_0_ and promoter p_*D*_. Substituting (21) into (20), the free dCas9 concentration dynamics can be written as:

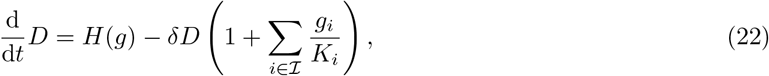

where

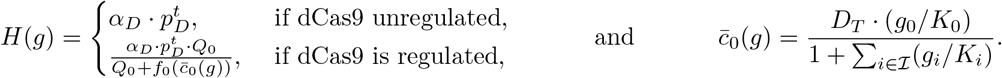

The total concentration of dCas9 protein *D_T_* is the summation of the concentration of apo-dCas9 *D* and the concentration of dCas9 proteins bound to sgRNAs:

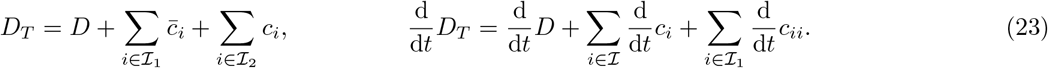

Hence, combining equations (5c)-(5e), and 20, we have the total dCas9 concentration dynamics:

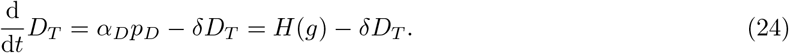

##### Dynamics of other proteins

The circuit produces a set of proteins other than dCas9. Their production rates may depend on CRISPRi-based regulation. These proteins are denoted by y_*i*_ with index *i* taking values in set 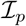. The synthesis and dilution of protein y_*i*_ are governed by the following chemical reactions:

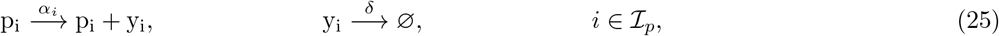

where *α_i_* is the synthesis rate constant of y_*i*_ from each copy of free promoter p_*i*_ and *δ* is the dilution rate constant. Hence, by mass action kinetics, the dynamics of *y_i_* can be written as

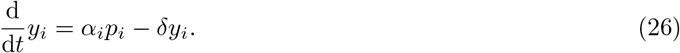

Transcription of y_*i*_ may be repressed by an sgRNA *g_q_*. In particular, *c_q_* (i.e., dCas9-sgRNA complex) may bind with p_*i*_ to form complex *c_qq_*, prohibiting transcription. Following (7), the QSS concentration of *c_qq_* is *c_qq_* = *p_i_c_q_*/*K_q_*. Since the copy number of DNA p_*i*_ is conserved, we have

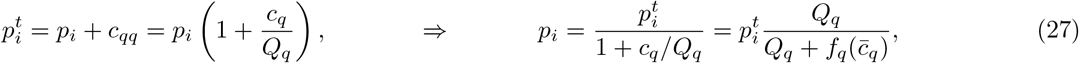

where 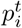 is the total concentration of the promoter driving protein y_*i*_ expression and 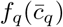 is defined as in (15). Substituting (27) into (26), the protein y_*i*_ dynamics can be written as:

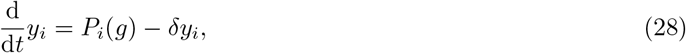

where

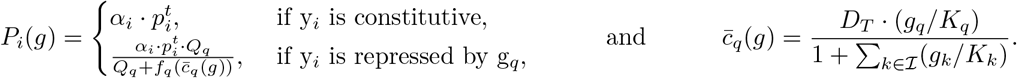

**Supplementary Table 7:** State variables and parameters invovled in the general CRISPRi-based circuit model (17), (24), and (28).

| State variables |  |
| --- | --- |
| $c_i$ | concentration of dCas9-$g_i$ complex |
| $c_{ii}$ | concentration of dCas9-$g_i$-promoter complex |
| $g_i$ | concentration of sgRNA $g_i$ |
| $D$ | concentration of apo-dCas9 protein |
| $D_T$ | concentration of total dCas9 protein |
| $y_i$ | concentration of protein $i$ |
| Parameters |  |
| $p_i^t$ | total promoter concentration driving $g_i$ or $y_i$ production |
| $p_D^t$ | total promoter concentration driving dCas9 protein production |
| $u_i$ | sgRNA $g_i$ synthesis rate constant from a single promoter |
| $\alpha_i$ | protein $y_i$ synthesis rate constant from a single promoter |
| $\alpha_D$ | dCas9 synthesis rate constant from a single promoter |
| $K_i$ | dissociation constant for sgRNA $g_i$ and dCas9 binding |
| $Q_i$ | dissociation constant for dCas9-$g_i$ complex and target DNA binding |
| $\delta$ | dilution due to cell fission |
| $\theta_i$ | sgRNA $g_i$ degradation rate constant |

##### Summary

Supplementary Table 7 summarizes the state variables and parameters in the general CRISPRi-based circuit model in (17), (24), and (28). Based on these equations, the effects of dCas9 competition manifest in the following way. If g_*j*_ (or y_*j*_) production is repressed by an sgRNA g_*i*_(*i* ≠ *j*), then the production rate of g_*j*_ (or y_*j*_) is not only dependent on *g_i_*, but also on the concentrations of all sgRNAs *g* in the circuit. This is because both functions *G_i_*(*g*) and *P_i_*(*g*) in (17) and (28) describing the production rates are *g*-dependent. To mitigate these unintended couplings arising from dCas9 competition, it is sufficient to maintain a constant level of apo-dCas9 concentration (*D*) that is independent of the concentration of competitor sgRNAs. Specifically, if *D* is a state-independent constant, then the concentrations of 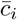 and *c_i_* in (7) depend only on *g_i_*. As a consequence, *p_j_* in (13) also depends only on *g_i_*, and the sgRNA dynamic model can be re-written as:

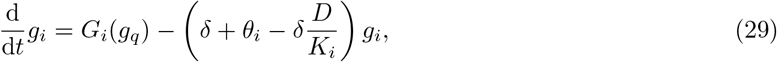

where

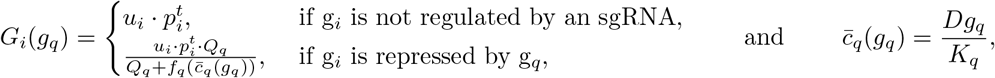

which does not depend on sgRNAs other than the intended regulator *g_q_*. Similarly, when apo-dCas9 level (*D*) is constant, the protein dynamics in (28) can be shown to depend only on the concentration of sgRNA repressing its promoter. We will show in Supplementary Note 4.3 that a practically constant *D* level can be achieved with the regulated dCas9 generator when the production rate of g_0_, which represses dCas9 production, is sufficiently high.

#### Supplementary Note 4.2 Models of the NOT gate and the cascade

Here, we apply the general modeling framework developed in Supplementary Note 4.1 to the NOT gate and the cascade. Since dCas9 binds to a tract of the sgRNA that is the same for all the guides (tracr-region) and since the 20bp annealing with the target have been designed to achieve the maximum repression efficiency, we assume throughout this section that the dissociation constants between sgRNAs and dCas9 protein are identical for all sgRNAs (i.e., *K_i_* = *K*), that the dissociation constant between dCas9-sgRNA complex with their target promoters are identical (i.e., *Q_i_* = *Q*), and that all sgRNAs have the same degradation rate constants (i.e., *θ_i_* = *θ*). The main outcomes do not depend on these assumptions. Because of these assumptions, we can write *f_i_*(·) = *f*(·) for the function defined in (15).

##### NOT gate with unregulated dCas9 generator

The CRISPRi-based NOT gate in Figure 1 consists of two sgRNAs g_1_ and g_2_. sgRNA g_1_ represses expression of the output protein RFP (y = y_4_) and sgRNA g_2_ does not have a DNA targeting site. Hence, we have 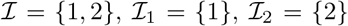, and 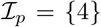. Since the transcription of g_1_ and g_2_ are not regulated by other sgRNAs, using (17), their dynamics are:

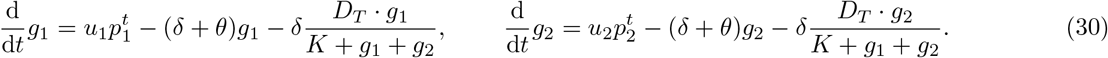

To model the fact that transcription of g_1_ is HSL-inducible, the transcription rate of g_1_ is modeled as a Hill function of HSL concentration: *u*_1_ = *u*_1_(HSL). Mathematical expression of this Hill function can be found in equation (52) in Supplementary Note 4.4. RFP expression is repressed by g_1_, hence, according to (28), we have

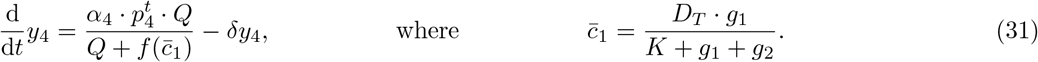

The total dCas9 concentration (*D_T_*) dynamics follow (24) with 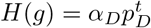, giving rise to

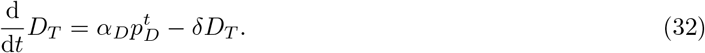

##### NOT gate with regulated dCas9 generator

For the NOT gate with regulated dCas9 generator, the circuit contains three sgRNA species, including g_0_ that represses dCas9 expression. Hence, 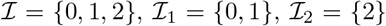, and 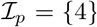. By (17), since g_1_ and g_2_ transcriptions are not regulated by other sgRNAs, we have

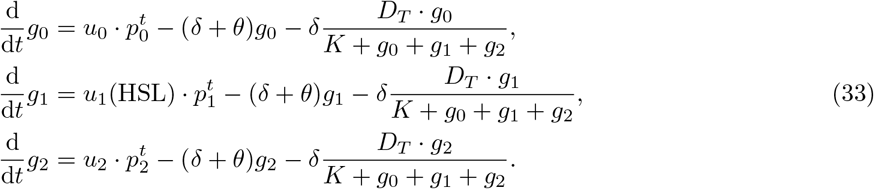

According to (28), RFP expression dynamics follow:

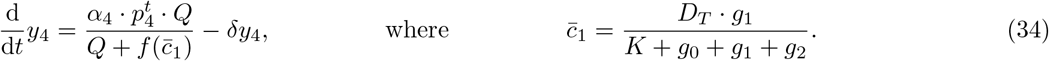

The dynamics of dCas9 protein are regulated and follow (24), giving rise to

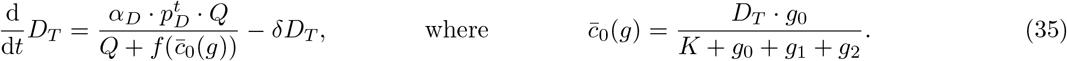

##### Cascade with unregulated dCas9 generator

The CRISPRi-based cascade shown in Figure 2 contains three sgRNAs. HSL-inducible g_1_ represses transcription of sgRNA g_3_, which subsequently represses expression of RFP y_4_. sgRNA g_2_ is transcribed constitutively as a resource competitor. Hence, we have 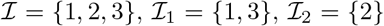, and 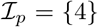. According to (17), the sgRNA dynamics follow:

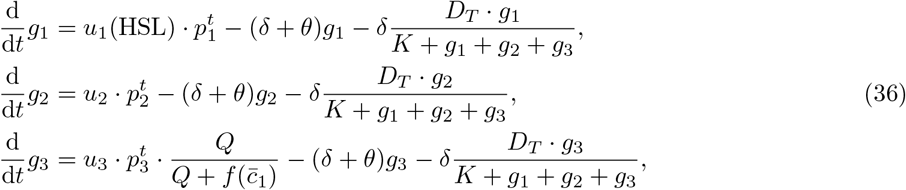

where

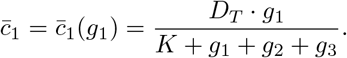

The expression of RFP is repressed by g_3_, hence, by (28), we have

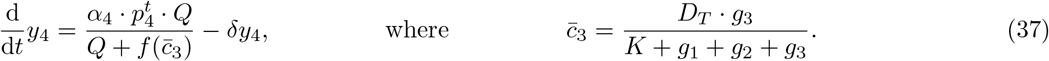

The total dCas9 concentration (*D_T_*) dynamics are unregulated and follow (24), giving rise to

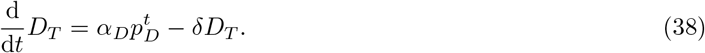

##### Cascade with regulated dCas9 generator

For the cascade circuit with regulated dCas9 generator, we take into account the additional sgRNA g_0_ to repress expression of dCas9. Hence, in this system, we have 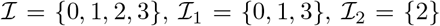, and 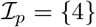. By (17), the sgRNA dynamics are:

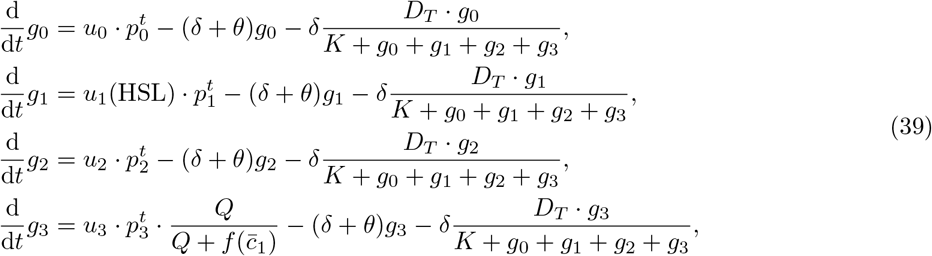

where

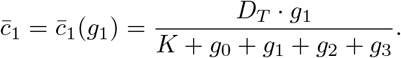

According to (28), the dynamics of RFP expression can be written as:

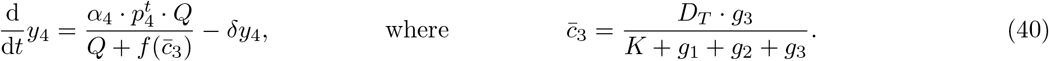

The total dCas9 concentration (*D_T_*) dynamics are regulated and follow (24), giving rise to:

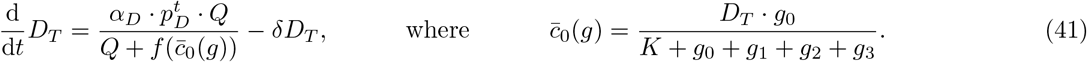

#### Supplementary Note 4.3 Model guided design of the regulated dCas9 generator

In this section, we consider the regulated dCas9 generator and demonstrate that increasing sufficiently the synthesis rate constant of g_0_ (i.e., *u*_0_) increases the robustness of a CRISPRi-NOT gate to the presence of a competitor sgRNA (i.e., g_2_). In particular, our analysis indicates that the sensitivity of apo-dCas9 concentration (*D*) to competitor sgRNA DNA copy number 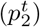 can be made arbitrarily small by increasing *u*_0_.

##### Sensitivity of apo-dCas9 concentration to competitor sgRNA

To be consistent with our notation in the main text, in addition to g_2_, sgRNA g_0_ represses dCas9 expression and g_1_ represses the output. The free sgRNA concentrations *g_i_* (*i* = 0,1, 2) depends on *D*. Specifically, at steady state, by setting the time derivative in (17) to zero, for *i* = 0,1, 2, we obtain:

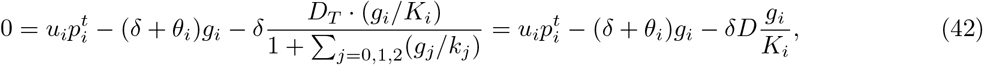

where we use 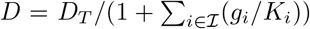 in (10) to attain the last equality. By (42), the steady state *g_i_* satisfies:

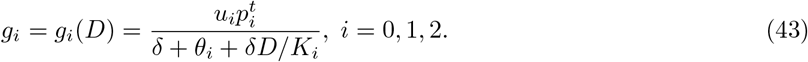

Since we study the system’s performance with 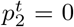 and 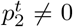, we specifically write 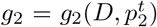. By setting the time derivative in (20) to zero, the steady state concentration of *D* can be solved from:

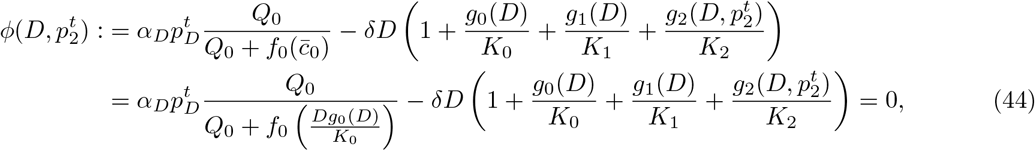

where we use 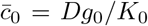 dervied in (7) in the last equality. The relative sensitivity of *D* to 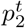 for the regulated dCas9 generator, which we denote by 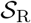, can be computed from (44) as:

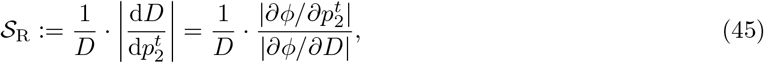

where we use the equality

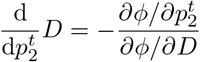

according to the implicit function theorem [S17]. From (43) and (44), we find

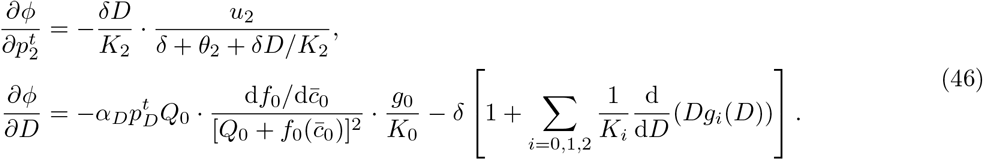

From (43), for *i* = 0,1, 2, we obtain

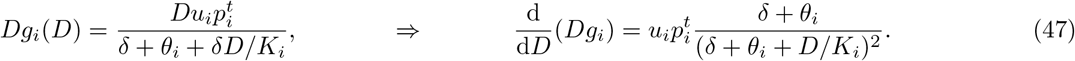

Substituting 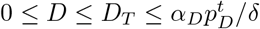 into (46) and (47), we find

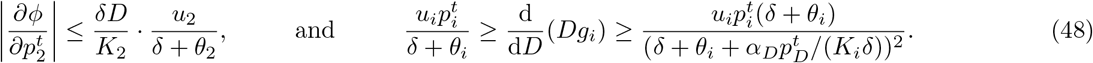

On the other hand, by the definition of *f*_0_ in (15), we have 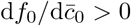 for all 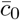. Combining this fact with the inequality in (48), we can find a lower bound for |*∂ϕ/∂D*| in (46):

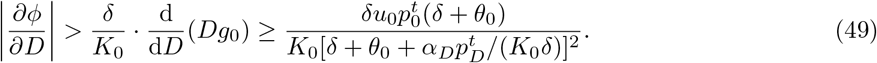

Substituting (48) and (49) into (45), and suppose that 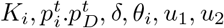, and *α_D_* are all positive constants, we can find an upper bound for 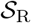 that depends on *u*_0_:

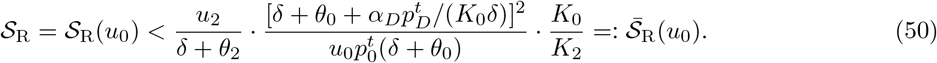

According to (50), the sensitivity upper bound 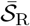 is a monotonically decreasing function of the g_0_ production rate constant *u*_0_. Additionally, 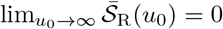. Because 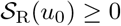 and 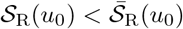, (50) implies that 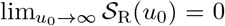. Hence, for a regulated NOT gate, if *u*_0_ is sufficiently large, then the apo-dCas9 concentration *D* becomes insensitive to the presence of g_2_ DNA (i.e., 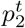). We verify this model prediction through simulations. In Supplementary Figure 6, we simulate the dose response curves of CRISPRi-based NOT gates with different g_0_ synthesis rate constants *u*_0_ and dCas9 protein synthesis rate constants *α_D_*. We find that as shown by our analysis, for each fixed *α_D_*, the dose response curves becomes independent of g_2_ when *u*_0_ is sufficiently large.

Physically, this increase in robustness is due to the presence of g_0_ that creates negative feedback actions on free dCas9 (*D*) dynamics. In particular, according to (22):

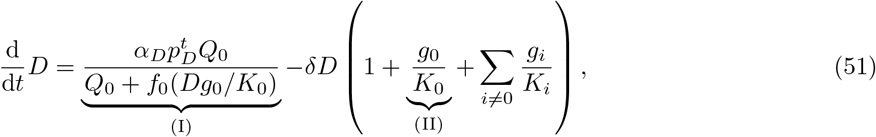

in which the feedback actions take two forms. On the one hand, with reference to the term labeled (I), a decrease in *D* leads to a decrease in *f*_0_(*Dg*_0_/*K*_0_) to increase the production rate of *D*. On the other hand, with reference to the term labeled (II) in equation, a drop in *D* also results in a decreased effective decay rate of *D*. Both forms of feedback actions contribute to the decrease in sensitivity of *D* to *g*_2_. Specifically, as we derive in (45) and (49), a small 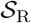 is due to a large |*∂ϕ/∂D*|, which is computed in (46). The feedback effect arising from dCas9 production rate change is manifested in the first term in (46), while the feedback effect arising from dCas9 effective decay rate change is manifested in the term encompassing d(*Dg*_0_)/d*D* in (46). Increasing the magnitude of both terms contributes to an increase in |*∂ϕ/∂D*|, hence, a decrease in 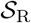. Therefore, both physical forms of feedback increase robustness of *D* to *g*_2_. As it can be observed from (51), increasing *g*_0_ (via, for example, increasing *u*_0_) can induce larger effects from both forms of feedback, which is consistent with our analysis in (50).

##### Experimental validation of sensitivity analysis

We designed an experiment to verify that sufficiently increasing *u*_0_ decreases 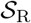. In particular, as shown in Supplementary Figure 7 (left and middle panels), we tested and compared two regulated dCas9 generators with same promoters and RBS for dCas9 production but with different promoters driving g_0_ production, which give rise to different *u*_0_ parameters. The two regulated dCas9 generators were co-transformed with the CRISPRi-NOT gate into *E. coli* NEB10B strain. The NOT gate either contains no competitor sgRNA or a competitor sgRNA g_2_ driven by the BBa_J23100 promoter. For the regulated dCas9 generator with g_0_ driven by the weaker BBa_J23116 promoter, the dose-response curve of the NOT gate is highly sensitive to the presence of g_2_. In fact, with reference to Supplementary Figure 7b (left panel), for low HSL levels, the fold change in RFP expression due to g_2_ production is similar to that induced by g_2_ when dCas9 production is unregulated (Figure 1c). On the other hand, with reference to Supplementary Figure 7b (middle panel), when the dCas9 generator is regulated by g_0_ driven by the stronger P108 promoter, RFP expression becomes insensitive to g_2_, indicating that robustness of apo-dCas9 concentration to competitor sgRNA production can indeed be achieved when the synthesis rate constant of g_0_ is sufficiently high.

##### Increasing dCas9 synthesis rate constant to maintain fold repression

Increasing g_0_ synthesis rate constant also increases repression on dCas9 synthesis, resulting in reduced total dCas9 concentration, hence reducing the concentration of dCas9-sgRNA complexes to repress target promoters. In fact, with reference to Supplementary Figure 6, for regulated dCas9 generators with small dCas9 synthesis rates (*α_D_*), increasing *u*_0_ leads to significant reductions in the fold repression of the NOT gates’ outputs. In order to achieve robustness to competitor sgRNA while maintaining fold repression by the gates’ sgRNAs, one can increase dCas9 synthesis rate in the regulated generator. This can be achieved by increasing the promoter strength driving the expression of dCas9 and/or the RBS strength of dCas9 transcript. In Supplementary Figure 6, our simulations indicate that when *u*_0_ is large, increasing dCas9 synthesis rate constant *α_D_* does not decrease robustness to competitor sgRNA. This result is further supported by our experiments in Supplementary Figure 7 (middle and right side panels). With reference to Supplementary Figure 7a, the regulated dCas9 generator in these two panels have identical *u*_0_ but different synthesis rate constants *α_D_* for dCas9 protein. In particular, while dCas9 expressions in both systems are driven by P104 promoters, the RBS strength of the system in the right panel is stronger than that of the system in the middle panel, indicting a substantial increase in *α_D_*. While in both systems, the competition effects by sgRNA g_2_ can be almost entirely mitigated by the regulator, the output of the system in the right panel shows larger fold repression. This difference is most significant when HSL=1 nM, where output level of the system with smaller *α_D_* is about 3x larger than that of the system with larger *α_D_*. Hence, based on these simulations and experiments, in order to increase robustness of CRISPRi-based circuits to dCas9 competition while maintaining similar fold repression, in the regulated dCas9 generator, we choose to transcribe g_0_ from a strong promoter (P112) while at the same time also express dCas9 protein from a stronger promoter (P104) using a stronger RBS.

**Supplementary Figure 6:**
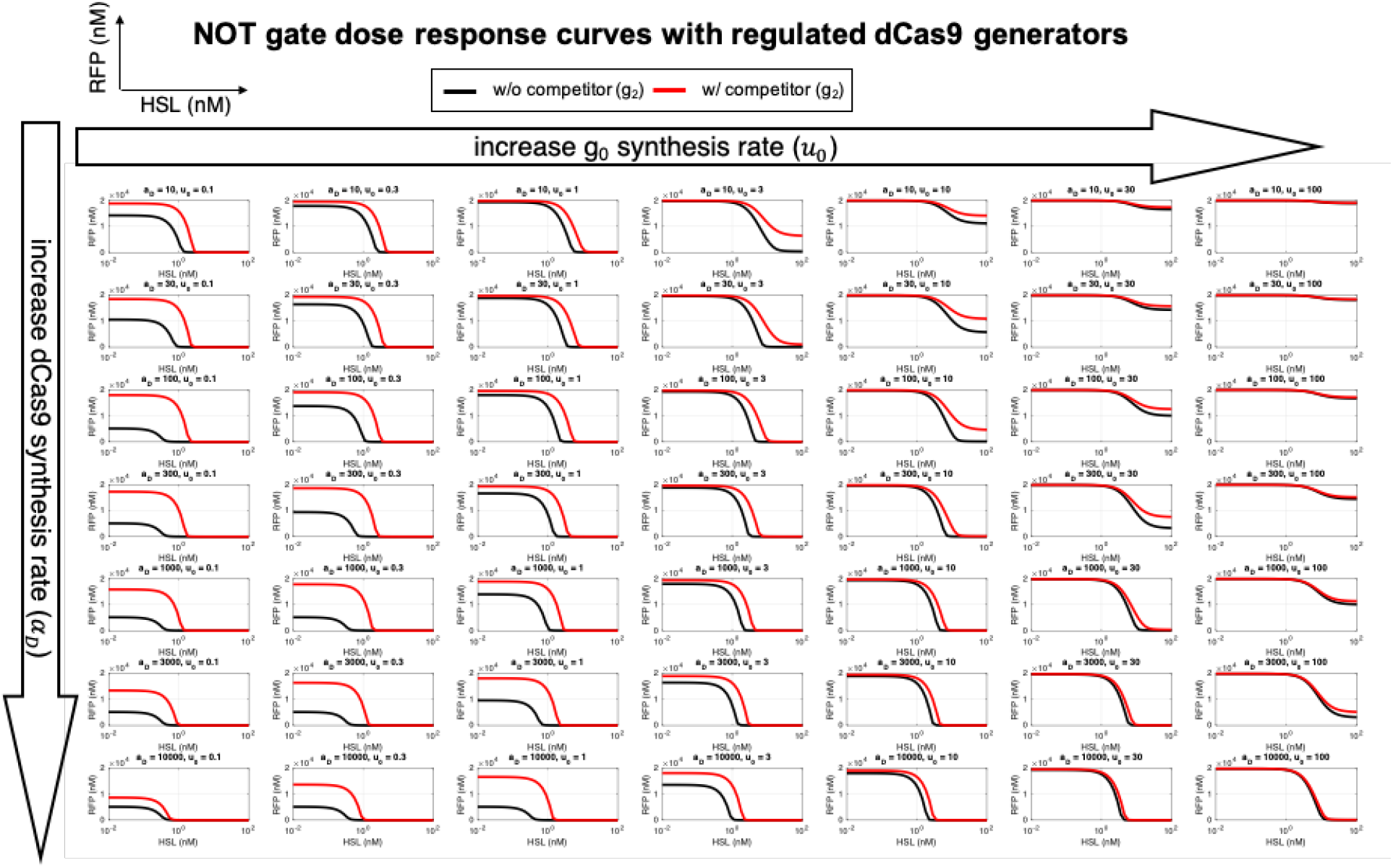
Dose response curves of the NOT gate with different regulated dCas9 generators. The generators have different dCas9 synthesis rate constants (*α_D_*) and g_0_ synthesis rate constants (*u*_0_). Black and red dose response curves correspond to systems without and with the competitor sgRNA g_2_, respectively. Model and parameters for simulations can be found in Supplementary Table 8 and Supplementary Table 9, respectively.

#### Supplementary Note 4.4 Numerical Simulations

Numerical simulations were carried out using MATLAB R2015b with variable step ODE solver `ode23s` to obtain the simulation results in Figures 1-2 and Supplementary Figure 6. In particular, the equations used for simulations are summarized in Supplementary Table 8.

**Supplementary Table 8:**
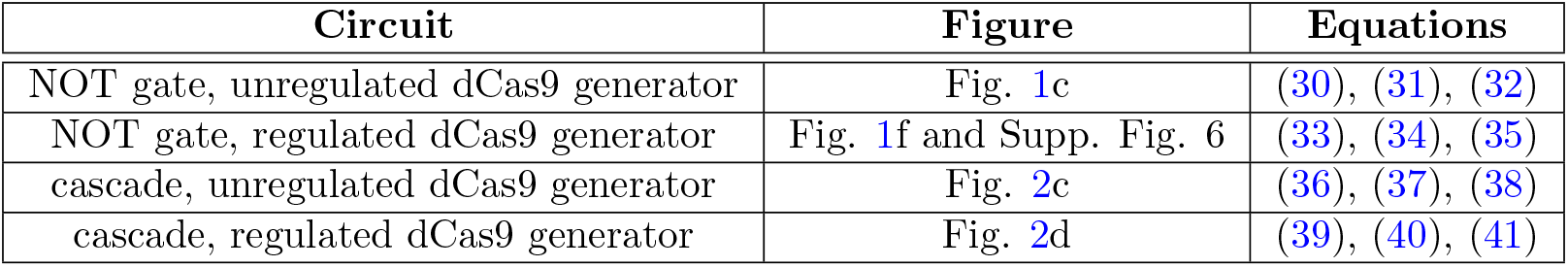
Equations used for simulations.

The parameters used for the simulations are listed in Supplementary Table 9. To obtain the plasmid concentrations, we follow the standard assumption that 1 copy/cell = 1 nM in bacteria *E. coli* [S18]. We use identical NOT gate parameters for simulations in the main text (Figure 1) and in Supplementary Figure 6.

**Supplementary Table 9:** Parameters used for simulations.

| Parameter | Unit | Fig. 1 | Fig. 2 | Supp. Fig. 6 | Ref |
| --- | --- | --- | --- | --- | --- |
| $u_2$ | [min<sup>-1</sup>] | 6 (J23116), 20 (J23100) | 40 (P108) | 20 | - |
| $u_3$ | | - | 24 | - | - |
| $u_0$ | | 65 | | varies | - |
| $\alpha_D$ | | 0.6 (unregulated), 6000 (regulated) | | varies | - |
| $\alpha_4$ | | 1 | | | - |
| $\delta$ | | 0.01 | | | experiment |
| $\theta$ | | 0.2 | | | [S12] |
| $p_0^t$ | [nM] | 30 | | | [S13] |
| $p_1^t = p_2^t$ | | 5 | 84 | 5 | |
| $p_3^t$ | | - | 84 | - | |
| $p_4^t$ | | 200 | 84 | 200 | |
| $p_D^t$ | | 30 | | | |
| $K$ | | 0.01 | | | [S14] |
| $Q$ | | 0.5 | | | [S15] |

The protein dilution rate constant *δ* is set to the average *E. coli* doubling time found in our experimental conditions. Since neither dCas9 nor RFP is targeted by a protease, we assume that protein degradation is negligible. The sgRNA synthesis rates *u_i_* for *i* = 0,1, 2, 3 are set to match the experimental qualitative I/O responses reported in Figure 1c, f and Figure 2c, d. The rank of the magnitudes of the synthesis rates matches the promoter strength rank in Supplementary Table 6. The increase in dCas9 synthesis rate in the regulated dCas9 generator reflects the design choice that dCas9 promoter and RBS are both much stronger in the regulated generator. Synthesis rate of *g*_1_ is modulated by the concentration of HSL. Hence, we use the following Hill function to model its synthesis rate *u*_1_:

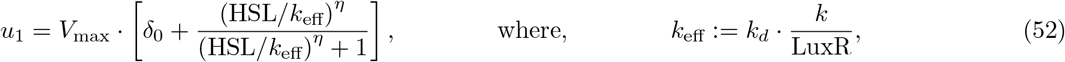

*V*_max_ is the maximum synthesis rate from the pLux promoter, *k* is the dissociation constant between HSL and LuxR protein, *k_d_* is the dissociation constant between HSL-LuxR complex and pLux promoter, *η* is the Hill coefficient, and *δ*_0_ represents the leakiness from the pLux promoter. We use the following parameters for equation (52) in simulations. The effective dissociation constant *k*_eff_ is smaller for the cascade system because LuxR protein is encoded on a higher copy plasmid there. Consequently, the concentration of LuxR is higher in the cascade system, leading to a reduced *k*_eff_.

**Supplementary Table 10:** Values of parameters describing Hill activation for g_1_ synthesis

| **Parameter** | $V_{max}$ [min<sup>-1</sup>] | $k_{\text{eff}}$ [nM] | $\eta$ | $\delta_0$ |
| --- | --- | --- | --- | --- |
| **NOT gate** | 50 | 8 | 2 | 0.006 |
| **Cascade** | 50 | 5 | 2 | 0.006 |

### Supplementary Note 5 Effect of sgRNA g0 transcription level on robustness of regulated dCas9 generators

To demonstrate the effect of sgRNA g0 transcription level on robustness of regulated dCas9 generators, we prepared two regulated dCas9 generators capable of transcribing different amounts of sgRNA g0. Specifically, we used the weak BBa_J23116 and the strong P112 promoters to transcribe sgRNA g0 of the regulated dCas9 generators in the pdCas9_CL2 and pdCas9_CL7 plasmids, respectively (Supplementary Table 11). The ribosome binding sites of *dCas9* genes in both plasmids are the BBa_B003J RBS. The pdCas9_CL2 and pdCas9_CL7 plasmids were co-transformed with the same plasmids of the plasmids 2 and 3 used in Supplementary Table 3 to create the constructs pCL92, pCL 94, pCL96 and pCL98 (see Supplementary Table 11) in order to compare to the constructs pCL87 and pCL89 (see Supplementary Table 3), respectively. The genetic diagrams of pCL96 and pCL98 are shown in Supplementary Figure 7a (the left and middle panels, respectively). We aim to compare how the dose-response curves of the CRISPRi-based NOT gate will change in the absence and the presence of the competitor sgRNA when apo-dCas9 proteins are expressed from the regulated dCas9 generator which transcribes sgRNA g0 in either low or high level.

Indeed, the response of the NOT gate is significantly affected by the competitor sgRNA at low HSL induction levels when the regulated dCas9 generator transcribes sgRNA g0 with the weak BBa_J23116 promoter as shown in Supplementary Figure 7b (the left panel). The fold-change can be up to 2-fold at the given level of the competitor sgRNA (Supplementary Figure 7c (the left panel)). This extent of the foldchange is similar to the one observed with the unregulated dCas9 generator (Figure 1c), suggesting that this regulated dCas9 generator dose not mitigate dCas9 competition because of low sgRNA g0 transcription. On the contrary, the response of the NOT gate is independent of the presence of the competitor sgRNA when the regulated dCas9 generator transcribes sgRNA g0 with the strong P112 promoter as shown in Supplementary Figure 7b (the middle panel). The fold-change remains practically unity (Supplementary Figure 7c (the middle panel)). The specific growth rates are similar among different constructs across the induced HSL concentrations as 0, 0.001, 0.01, 0.1, 1, and 50 nM (Supplementary Figure 7d, the left and middle panels). The experimental data are in agreement with the sensitivity analysis of the regulated dCas9 generator (Supplementary Note 4.3) such that increasing the synthesis rate of sgRNA g0 increases the robustness of a CRISPRi-based NOT gate to the presence of a competitor sgRNA.

#### Effect of dCas9 production rate on fold repression of CRISPRi-based NOT gate

When the ribosome binding site of *dCas9* gene is changed from the BBa_B0034 to the RBS1, comparing the genetic diagram of Supplementary Figure 7a in the middle panel to the one in the right-side panel, this change significantly increase dCas9 production rate in the regulated dCas9 generator. The fold-changes remain almost the same as shown in the middle and right-side panels of Supplementary Figure 7c, but the dose-response curves, especially at 1 nM HSL, show a better repression on the CRISPRi target in the right panel than in the middle panel of Supplementary Figure 7b. This suggests that higher dCas9 production rate does not affect the robustness of the regulated dCas9 generator but improves fold repression of the CRISPRi-based NOT gate. Similar specific growth rates of different constructs across the induced HSL concentrations, as shown in Supplementary Figure 7d, suggest that increased dCas9 level did not lead to any cytotoxicity. The experimental data are in agreement with the analysis of Supplementary Note 4.3 and the simulations of Supplementary Figure 6, according to which the production rate of dCas9 can be increased to improve the fold repression of CRISPRi while keeping robustness if the sgRNA g0 production rate is sufficiently large.

**Supplementary Table 11:** List of the constructs used in Supplementary Figure 7. Each construct is obtained from co-transforming the indicated plasmids into *E. coli* NEB10B strain.

| construct | plasmid 1 (Kan) | plasmid 2 (Amp)<sup>10</sup> | plasmid 3 (Cm) | sgRNA g0's promoter | sgRNA g2's promoter |
| --- | --- | --- | --- | --- | --- |
| pCL92 | pdCas9_CL2 | I13521-target | AEgPtet_No_competitor | BBa_J23116 | cassette absent |
| pCL94 | pdCas9_CL7 | I13521-target | AEgPtet_No_competitor | P112 | cassette absent |
| pCL96 | pdCas9_CL2 | I13521-target | AEgPtet_100gPlac | BBa_J23116 | BBa_J23100 |
| pCL98 | pdCas9_CL7 | I13521-target | AEgPtet_100gPlac | P112 | BBa_J23100 |

**Supplementary Figure 7:**
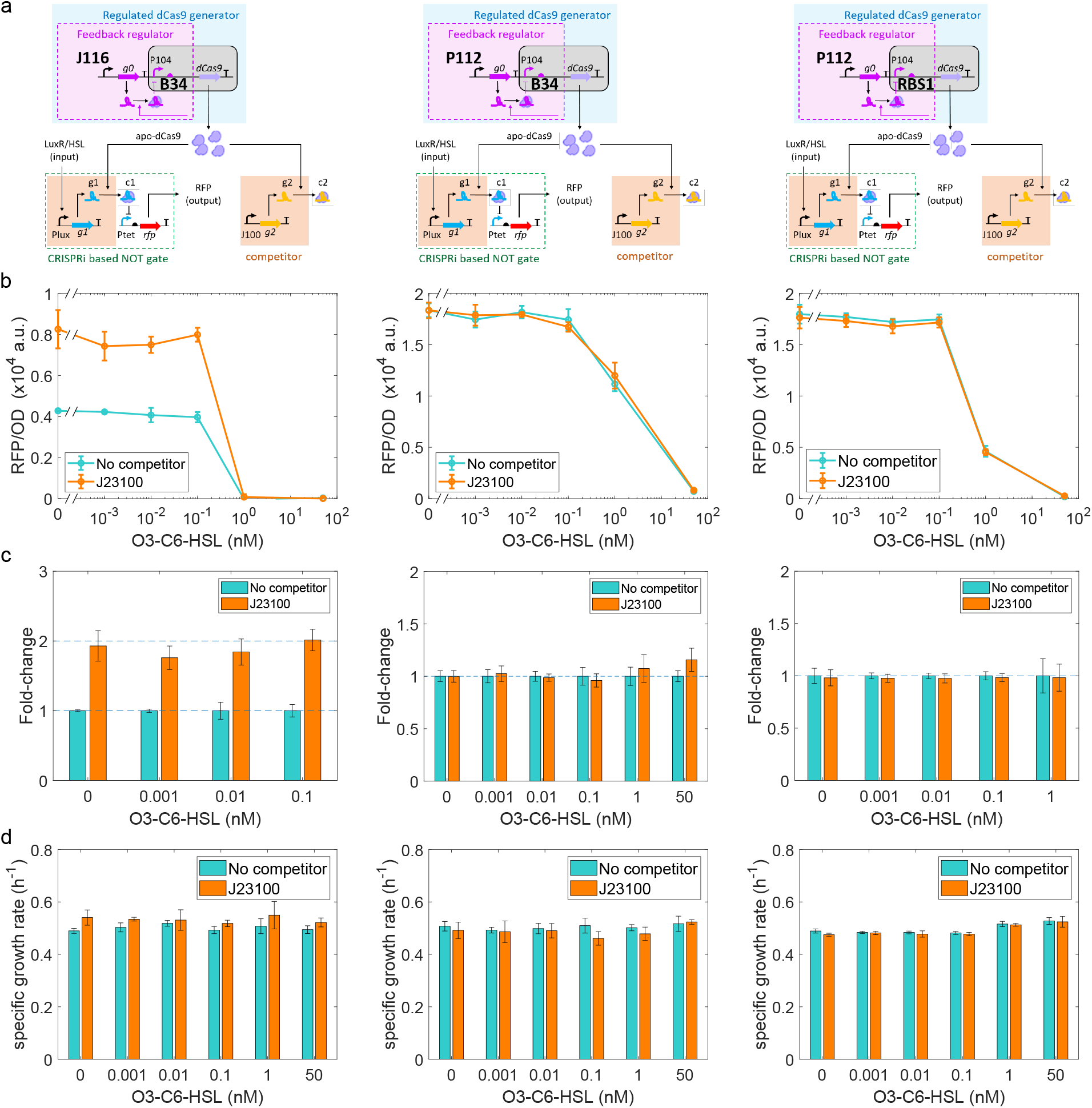
Abundant sgRNA g0 contributes to robustness of regulated dCas9 generator and higher dCas9 production rate improves fold repression. Apo-dCas9 proteins are expressed by the regulated dCas9 generator in which the promoter of sgRNA g0 and the ribosome binding site of *dCas9* gene are in a pair as (promoter, RBS) as (BBa_J23116, BBa_B0034), (P112, BBa_B0034), and (P112, RBS1) in the left, middle, and right-side panels, respectively. (a) Genetic circuit of the CRISPRi-based NOT gate and the competitor module is identical to the one used in Figure 1. The competitor module is either absent or using the BBa_J23100 promoter to transcribe the competitor sgRNA g2. (b) Comparison of dose-response curves in the absence or presence of the competitor module. (c) Fold-changes at a given HSL induction were computed, according to Online Methods Equation 1, by dividing the RFP/OD value of a construct by the one of the construct lacking the competitor module. (d) Specific growth rate of each construct at a given induced condition. The culture of *E. coli* NEB10B cells grew at 30 °C in M9 medium. Data with error bars represent mean values with standard deviations from three experimental repeats by microplate photometer. The data in all right-side panels are reproduced from the data in Figure 1 and Supplementary Figure 4.

### Supplementary Note 6 Regulator to neutralize dCas9 competition in other CRISPRi-based NOT-gate circuits, growth conditions, and strains

To demonstrate the effect of dCas9 competition on CRISPRi-based NOT gates in different contexts and to verify the ability of the regulated dCas9 generator to mitigate competition, we varied DNA copy number of NOT gate and competitor, *E. coli* strain, and the NOT gate input regulator molecule from the ones of the circuit in Figure 1.

First, we investigated the extent of competition and the ability of the dCas9 generator to mitigate competition when a CRISPRi-based NOT gate and a competitor sgRNA are both expressed by a plasmid with higher copy numbers. We created the plasmid pHH41 in which the origin of replication is pSC101(E93G), which was reported to be ~84 copies/cell [S2]. As a comparison, the origin of replication of AEgPtet plasmids used in Figure 1 is pSC101, which was reported to be ~5 copies/cell [S10]. The pHH41 plasmid encodes both the NOT gate and the competitor sgRNA cassette. When we use the BBa_J23116 promoter to transcribe the competitor sgRNA, we call this plasmid pHH41_1 (Supplementary Figure 8). When instead we use the BBa_J23100, pTrc, and BBa_J23119 promoters to transcribe the competitor sgRNA, we call the plasmids pHH41_2, pHH41_3, and pHH41_4, respectively, as listed in Supplementary Table 12. Much stronger constitutive promoters such as the pTrc and BBa_J23119 promoters [S4] were selected to reach the respective higher concentrations of the competitor sgRNA. The competitor sgRNA cassette is located in the intergenic region between the RFP expression cassette and the LuxR expression cassette. The targeting sgRNA of the NOT gate represses the strong P105 promoter of the *mRFP1* gene and is controlled by the plux promoter and transcription factor LuxR. The guide sequence of the targeting sgRNA is labeled as sgP105 (Supplementary Table 2). The pHH41 plasmids were transformed concurrently with either the pdCas9_OP or the pdCas9_CL plasmid into *E. coli* TOP10. The growth condition is at 30 °C and using glucose as the carbon source in M9610 medium. M9610 medium is buffered at pH 6 [S11].

The circuit diagrams of the constructs pOP4, pOP5, pOP6, and pOP7 are shown in Supplementary Figure 9a, where the Pc promoter is BBa_J23116, BBa_J23100, BBa_J23119, and pTrc promoter, respectively. These constructs use unregulated dCas9 generator. LuxR’s effector O3-C6-HSL induces the NOT gate to repress RFP expression in the presence of different amounts of the competitor sgRNA. Dose-response curves are shown in Supplementary Figure 9b. The more the competitor sgRNA is transcribed by a stronger promoter, the more the shape of a dose-response curve deviates from the one of the curve of the pOP4 construct. Foldchanges were not computed when the concentration of HSL is higher than 30 nM because in such induced conditions, the RFP/OD values of pOP4 are approximately zero and the fold-changes become undefined. The maximal fold-change observed was up to 25-fold at 1 nM HSL when the competitor sgRNA is transcribed by the pTrc promoter. Specific growth rates at steady state were not affected across different induced conditions (Supplementary Figure 9d).

When using the regulated dCas9 generator, the constructs pCL28, pCL29, pCL30, and pCL31 should be compared to the constructs pOP4, pOP5, pOP6, and pOP7, respectively. The genetic diagram is shown in Supplementary Figure 9e. Dose-response curves of these constructs, remain practically the same even when the competitor sgRNA is transcribed by the strong pTrc promoter (Supplementary Figure 9f). The definable fold-changes remain almost equal (Supplementary Figure 9g). Comparing to the fold-changes in Supplementary Figure 9c, the regulated dCas9 generator can neutralize dCas9 competition even at higher copy numbers of the NOT gate and the competitor. Specific growth rates at steady state were barely affected across different induced conditions (Supplementary Figure 9h) and were slightly lower than the respective ones in Supplementary Figure 9d.

We next investigated whether the extent of competition and the ability of the regulated dCas9 generator to mitigate it could generalize to the use of other input regulators for the NOT gate, repressors, specifically. We thus changed the transcriptional regulator from LuxR activator to TetR repressor and the cognate promoter from plux promoter to Ptet promoter. Thus, the pHH41 plasmids were modified accordingly into what we called the pHH43 plasmids (Supplementary Figure 8). In parallel with the constructions of pOP4-pOP7 and pCL28-pCL31 constructs, we created pOP9-pOP12 and pCL33-36 constructs with the component plasmids listed in Supplementary Table 12. They were characterized in the same growth condition as pOP4-pOP7 and pCL28-pCL31 constructs.

The circuit diagrams of the constructs pOP9, pOP10, pOP11, and pOP12 are shown in Supplementary Figure 10a, where the Pc promoter is one among BBa_J23116, BBa_J23100, BBa_J23119, and pTrc, respectively. These constructs use the unregulated dCas9 generator. TetR’s effector anhydrotetracycline (aTc) induces the NOT gate to repress RFP expression in the presence of different amounts of the competitor sgRNA. Dose-response curves are shown in Supplementary Figure 10b. We could observe substantial dCas9 competition in such context, indicating that the effect of dCas9 competition is independent of the type of the regulation of the NOT gate. Fold-changes were computed as explained in Methods. The maximal fold-change up to 13-fold can be observed at 100 nM aTc when the competitor sgRNA is transcribed by the pTrc promoter. Inappreciable changes in specific growth rates at steady state were observed across different induced conditions (Supplementary Figure 10d).

When using the regulated dCas9 generator, the constructs pCL33, pCL34, pCL35, and pCL36 were built as a comparison to the constructs pOP9, pOP10, pOP11 and pOP12, respectively. The genetic diagrams are shown in Supplementary Figure 10e. From dose-response curves (Supplementary Figure 10f) and fold-changes at a given induced condition (Supplementary Figure 10g), we observed consistent results supporting that the regulated dCas9 generator can neutralize dCas9 competition in the current genetic context and growth condition. Specific growth rates at steady state were scarcely affected across different induced conditions (i.e. 0, 30, 60, 100, 300 nM aTc, Supplementary Figure 10h).

**Supplementary Figure 8:**
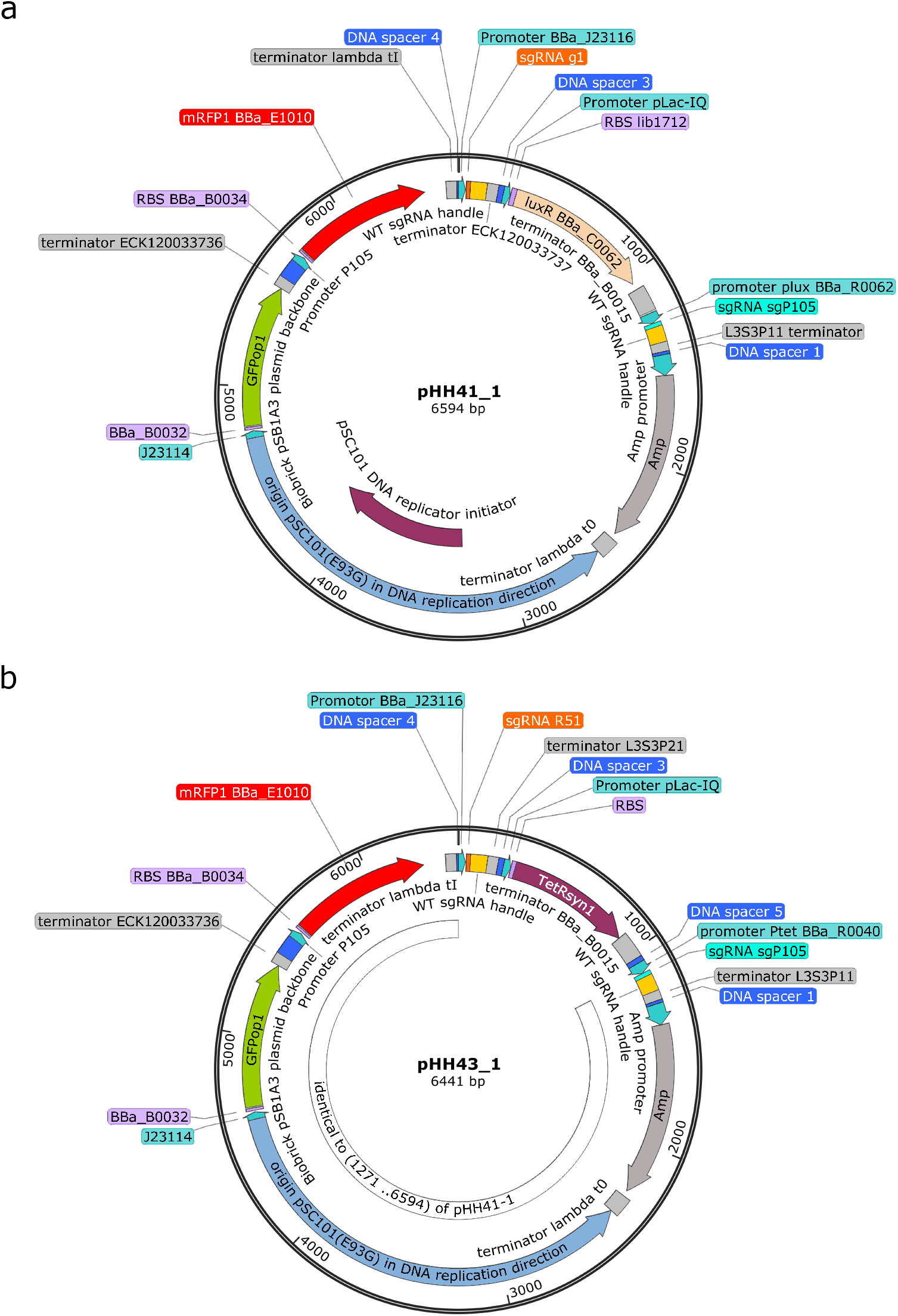
Maps of the pHH41_1 and pHH43_1 plasmids which were used in the constructs listed in Supplementary Table 12. (a) pHH41_1 plasmid encodes a CRISPRi-based NOT gate which is regulated by transcriptional activator LuxR and the competitor sgRNA which is transcribed by BBa_J23116 promoter. The targeting sgRNA of the NOT gate and the competitor sgRNA are on map position 1271 and 40, respectively. (b) pHH43_1 plasmid encodes a CRISPRi-based NOT gate which is regulated by transcriptional repressor TetR and the competitor sgRNA which is transcribed by BBa_J23116 promoter. The targeting sgRNA of the NOT gate and the competitor sgRNA are on map position 1118 and 40, respectively. The derivation of pHH41 and pHH43 series plasmids are detailed in Supplementary Note 6. The guide sequences of sgRNAs are listed in Supplementary Table 2.

**Supplementary Table 12:**
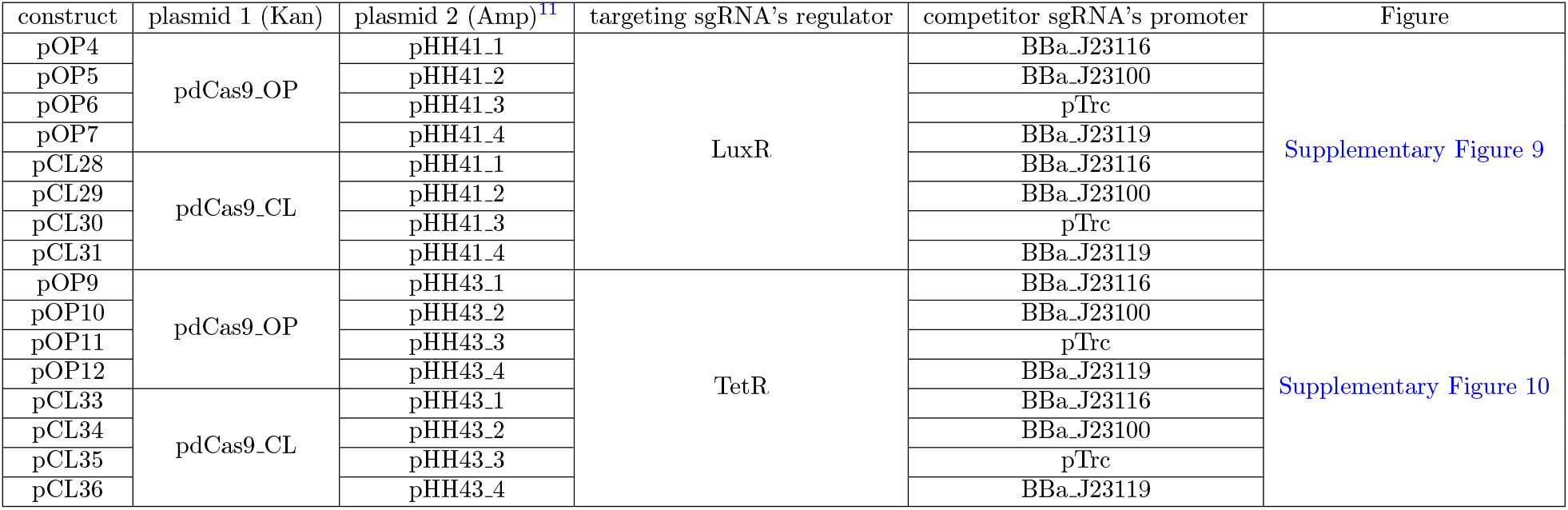
List of the constructs used in Supplementary Figure 9 and Supplementary Figure 10. Constructs were co-transformed with the indicated plasmid 1 and plasmid 2 into *E. coli* TOP10 strain. The regulator of the targeting sgRNA and the promoter of the competitor sgRNA are listed.

**Supplementary Figure 9:**
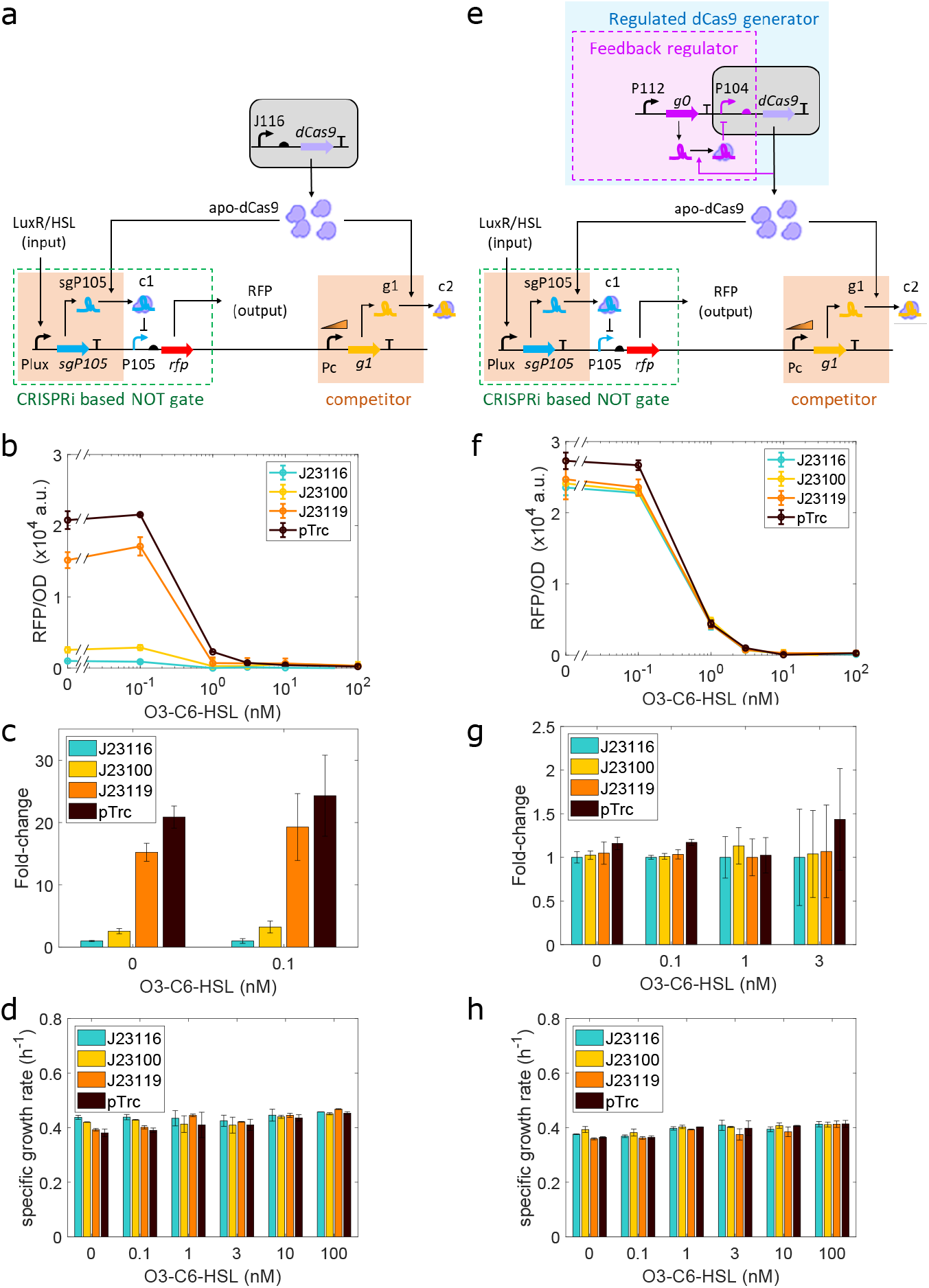
Neutralization of dCas9 competition in the high-copy number CRISPRi-based NOT gate. The panels (a-d) and (e-h) exhibit the results when using the unregulated and regulated dCas9 generator, respectively. (a) Genetic circuit of the CRISPRi-based NOT gate and the competitor sgRNA is encoded in a plasmid using pSC101(E93G) origin (~84 copies). The Pc promoter of the competitor module is BBa_J23116. BBa_J23100, BBa_J23119. or pTrc promoter to tune the level of the competitor sgRNA. Apo-dCas9 proteins are expressed from the unregulated dCas9 generator in a plasmid using p15A origin. (b) Comparison of dose-response curves in the presence of different amounts of the competitor sgRNA transcribed by the indicated promoter. (c) Fold-changes at a given HSL induction were computed, as in Online Methods Equation 1, by dividing the RFP/OD value of a construct by the one of the construct which uses the BBa_J23116 promoter to transcribe the competitor sgRNA. (d) Specific growth rates of each construct at a given induced condition. (e) Genetic circuit of the CRISPRi-based NOT gate and the competitor is the same as in (a). Apo-dCas9 proteins are expressed from the regulated dCas9 generator in a plasmid using p15A origin. (f) Comparison of dose-response curves in the presence of different amounts of the competitor sgRNA transcribed by the indicated promoter. (g) Fold-changes were computed in the same way as in (c). (h) Specific growth rates of each construct at a given induced condition. The culture of *E. coli* TOP10 cells grew at 30 °C in M9610 medium. Data with error bars represent mean values with standard deviations from two biological repeats by microplate photometer.

**Supplementary Figure 10:**
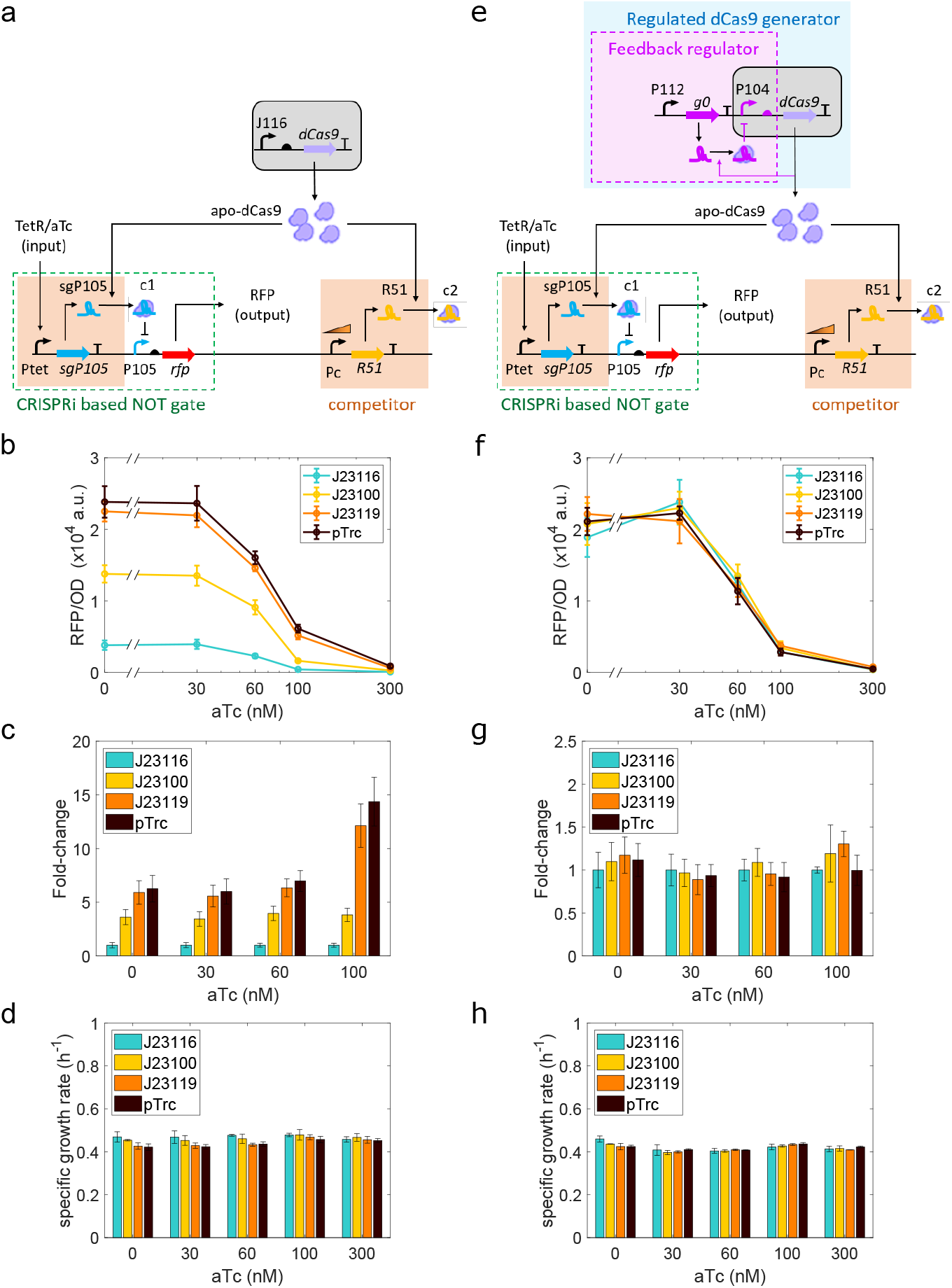
Neutralization of dCas9 competition in the high copy number CRISPRi-based NOT gate with input regulator TetR and its effector aTc. The panels (a-d) and (e-h) exhibit the results when using the unregulated and regulated dCas9 generator, respectively. (a) Genetic circuit of the CRISPRi-based NOT gate and the competitor sgRNA is encoded in a plasmid using pSC101(E93G) origin. The Pc promoter of the competitor module is the BBa_J23116, BBa_J23100, BBa_J23119, or pTrc promoter to tune the level of the competitor sgRNA. Apo-dCas9 proteins are expressed from the unregulated dCas9 generator in a plasmid using p15A origin. (b) Comparison of dose-response curves in the presence of different concentrations of the competitor sgRNA transcribed by the indicated promoter. (c) Fold-changes at a given aTc induction were computed, as in Online Methods Equation 1, by dividing the RFP/OD value of a construct by the one of the construct which uses the BBa_J23116 promoter to transcribe the competitor sgRNA. (d) Specific growth rates of each construct at a given induced condition. (e) Genetic circuit of the CRISPRi-based NOT gate and the competitor is the same as in (a). Apo-dCas9 proteins are expressed from the regulated dCas9 generator in a plasmid using p15A origin. (f) Comparison of dose-response curves in the presence of different concentrations of the competitor sgRNA transcribed by the indicated promoter. (g) Similarly, fold-changes were computed in the same way as in (c). (h) Specific growth rates of each construct at a given induced condition. The culture of *E. coli* TOP10 cells grew at 30 °C in M9610 medium. Data with error bars represent mean values with standard deviations from three biological repeats by microplate photometer.

## Footnotes

1 Kan, Amp, and Cm stand for kanamycin, ampicillin, and chloramphenicol, respectively.

2 Data are adopted from [S2]. n.a., not available.

3 The copy number of pUC19-derived pMB1 origin was reported as 500-700 from [S3] or 100-300 from the iGEM information on pSB1A2 plasmid from the Standard Registry of Biological parts.

4 The P075, P104, and P105 promoters are adopted from Ec-TTL-P075, P104, and P105 promoters [S4], respectively. Other promoters with a BBa_ number are adopted from iGEM registry.

5 For the competitor sgRNAs, the intended target sequence is absent in the indicated plasmids and host cell strain.

6 All plasmids of the plasmid groups 2 and 3 are from [S1].

7 The BBa_J23116 and BBa_J23100 promoters are from the iGEM Registry of Standard Biological Parts. Maps of the plasmids 1, 2, and 3 are shown in Supplementary Figure 1 and Supplementary Figure 2.

8 The pHH50-I plasmid encodes the competitor sgRNA g2 cassette (i.e. promoter, sgRNA, and terminator). The pHH50-IV plasmid keeps the P108 promoter and the synthetic terminator 4 of the cassette but sgRNA g2 is absent.

9 This Plux carries a deletion of the last 3 nucleotides (AAA) to set the transcription start site at +1; the deletion leads to a reduction of protein synthesis rate of ~3.7 times, as reported in [S1]

10 The plasmids 2 and 3 are from the same set of the plasmids 2 and 3 in Supplementary Table 3.

11 Maps of the pHH41_1 and pHH43_ 1 plasmids are shown in Supplementary Figure 8. Variants of pHH41 and pHH43 plasmids are only different in the promoter of the competitor sgRNA. The usage of these constructs is detailed in Supplementary Note 6.

